# Active Learning–Guided Peptide Design for Modulating Condensate Properties upon Recruitment

**DOI:** 10.64898/2026.06.06.730477

**Authors:** Kumar Gaurav, Rodrique Badr, Arya Changiarath, Vasileios A. Xenidis, Lukas S. Stelzl

## Abstract

The physical properties of biomolecular condensates, which form through phase separation, are central to their organisation and function and are increasingly implicated in disease mechanisms. To function correctly, condensates often recruit client biomolecules such as peptides and RNAs. This recruitment is not only essential to condensate biology but also offers an opportunity to engineer clients that tune the material properties of the condensate. Here we studied the previously characterised MUT-16 condensate, the scaffold of a membraneless organelle in *C. elegans* that recruits the N-terminal prion-like domain of MUT-8, a process essential for RNA silencing. We used coarse-grained molecular dynamics simulations to model and design peptide variants that interact with the scaffold protein MUT-16 and modulate its physical properties. To guide this design efficiently, we applied an active learning framework, which combines Bayesian optimization with neural networks to iteratively select the most informative peptide variants for simulation. This strategy reduced the number of simulations required while allowing the model to learn the sequence–property relationships that govern condensate behavior. Overall, this physics-based, learning-guided framework offered a computational approach to exploring how peptides can be engineered to influence condensate properties, and may inform the rational design of synthetic condensates.

## Introduction

Intrinsically disordered proteins and regions (here collectively termed IDPs) constitute a diverse class of proteins characterized by pronounced structural heterogeneity and a lack of stable folded conformations.^1,2^ Their distinct amino acid compositions prevent the formation of well-defined tertiary structures, and they are instead described by rapidly interconverting ensembles of conformations.^3,4^ The dynamic and disordered nature of IDPs underlies many of their biological functions, including acting as flexible linkers, spacers, and mediators of molecular recognition through short linear motifs.^5,6^ Owing to their multivalent interaction capacity, IDPs frequently participate in signaling and transcriptional regulation and play a central role in the spatial organization of cellular matter.^7,8^ In particular, IDPs can drive the formation of membraneless organelles via phase separation, resulting in the coexistence of a protein-rich dense phase and a dilute phase when the average concentration is above a characteristic saturation concentration (*c*_sat_).^8^ The molecular interactions underlying this process span both *homotypic* contacts (self-association of like molecules) and *heterotypic* contacts (association of different species), whose balance shapes the resulting dense phase.^9^ An interplay of different molecular interactions, often weak and multivalent drives phase separation.^10–13^ Such interactions on the molecular scale also determine the collective behaviour of molecules within the condensate. Previous studies have demonstrated that the dense phase often exhibits material properties similar to those of a viscoelastic material.^14–16^ The material properties of biomolecular condensates span a broad spectrum, ranging from highly dynamic, liquid-like assemblies to more rigid, gel-like or solid-like states. ^17^ These physical characteristics critically influence their biological functions, including processes associated with cellular ageing and responses to environmental or intracellular cues. ^18^ Importantly, alterations in condensate material properties have been strongly implicated in disease pathogenesis, particularly in neurodegenerative disorders^19^ and oncogenic pathologies, where aberrant transitions toward more solid-like states can disrupt normal cellular regulation.^20,21^

The material properties of biomolecular condensates, including viscosity, surface tension, viscoelasticity, and ageing, are encoded in the primary sequence of the intrinsically disordered proteins that undergo phase separation.^13,22,23^ In particular, residue identity, valency, and spatial patterning of interaction motifs determine the network of transient intermolecular interactions that collectively govern condensate thermodynamics and mesoscale physical behavior.^13^ This intimate coupling between protein sequence and condensate material properties enables rational protein design strategies to systematically tune and modulate the physical behavior of the resulting condensates. ^24–28^ In this context, computational design approaches coupled with molecular dynamics simulations provide an efficient and cost effective framework to explore the sequence space and predict material properties prior to experimental validation.

In addition to material properties, condensate function is strongly influenced by the interface between the dense and dilute phases.29–31 Previous studies have implicated condensate interfaces in fibril formation and liquid-to-solid transitions.32–34 The mesoscale properties of the interface are, in turn, determined by protein conformations at the interface, as captured by the radius of gyration (*R_g_*) and orientational order parameter (*S_z_*).29,35 Furthermore, von Bülow *et al.*35 demonstrated a strong correlation between the transfer free energy, Δ*G*_trans_, and interfacial conformational properties, particularly the swelling ratio *R_g,_*_interface_*/R_g,_*_dense_ and the orientational order parameter *S_z,_*_interface_.

Molecular dynamics simulations have become a central tool in understanding and engineering phase separating proteins. ^10,27,35–40^ Continuous improvements in molecular force fields enable increasingly accurate descriptions of intermolecular interactions, allowing simulations to reproduce and predict phase separation propensities across diverse sequences.^36,37,41^ Atomistic simulations provide mechanistic insight into how residue identity, valency, and sequence context give rise to emergent condensate properties.^10,42,43^ Complementarily, coarse-grained simulation models, including residue-level coarse-grained simulation models, enable access to larger system sizes and longer time scales, thereby facilitating the exploration of phase diagrams, collective effects, and emergent material properties at a reduced computational cost.^23,36,44–46^ However, establishing a quantitative and predictive sequence-property relationship that enables protein design without exhaustive simulations or experiments remains challenging, owing to the vastness of the sequence space and the strong context dependence of amino acid interactions.

Machine learning offers a powerful framework to address this challenge in the context of rational protein design of IDPs.^22,24,25,28,37,40,49–52^ Hybrid approaches that combine physics-based simulations with data-driven models have proven successful in learning the sequence-encoded determinants of phase behaviour,^22,38,53,54^ and recent work has demonstrated the de novo design of peptides that partition to condensate interfaces.^55^ In particular, large pre-trained protein language models trained on a myriad of natural sequences, generate high dimensional embeddings that capture biochemical, structural, and evolutionary information, thereby providing informative representations for downstream prediction tasks and efficient exploration of the sequence space.^48,56–60^ Because labeled data are often costly to obtain, active learning provides an iterative strategy in which a predictive model is trained on an initial dataset and subsequently queries the most informative new sequences for label-ing.^22,40,61,62^ Changiarath et al. demonstrated that active learning using featurisation with a protein language model and coarse-grained simulations can be used for designing IDR-IDR interactions and also IDR partitioning to condensates and ultimately to enables control of condensate morphology in simulations.^26^ By prioritizing sequences with the most desirable qualities, these approaches minimize the number of expensive simulations or experiments required. Together, physics-informed machine learning, protein language model embeddings, and uncertainty-aware active learning^47^ establish an effective approach for designing proteins with tailored condensate material properties.^22,63^

We apply this approach to MUT-16 condensates in the presence of short peptides. MUT-16 is a scaffolding protein that nucleates the formation of the membraneless organelles known as *Mutator foci* in *C. elegans*.^64^ These condensates are essential for the amplification of small interfering RNAs (siRNAs) during RNA interference (RNAi)–mediated gene silencing. RNAi is a conserved gene regulatory pathway that preserves genome integrity by suppressing viruses and transposable elements and is required for diverse biological processes, including gametogenesis, chromosome segregation, and development.^65,66^ The assembly of MUT-16 into *Mutator foci* and its ability to recruit key components of the siRNA amplification machinery—including the RNA-dependent RNA polymerase RRF-1, the nucleotidyltrans-ferase MUT-2, the DEAD-box RNA helicase MUT-14, the 3*′*–5*′* exoribonuclease MUT-7, and MUT-8^67,68^ (also known as RDE-2)—are therefore central to the proper execution of RNAi-mediated gene regulation.^69,70^ MUT-16 is predominantly intrinsically disordered, and its phase separation behavior is encoded within specific regions of its sequence. A segment spanning residues 773–945, termed the foci-forming region (FFR), is both necessary and sufficient for condensate formation, whereas a distinct MUT-8-binding region (M8BR, residues 633–772) mediates interaction with MUT-8.^64,71^ The combined M8BR+FFR construct undergoes liquid–liquid phase separation *in vitro*, forming liquid-like condensates that recapitulate key features of *Mutator foci*.^71^ Recruitment of MUT-8,whose N-terminus contains a prion-like domain and directly interacts with the M8BR of MUT-16, is essential for RNA silencing and suggests that heterotypic interactions play a critical role in regulating condensate composition and function.^64,71^ Previous work from our group demonstrated that Arg residues in MUT-16 and Tyr residues in MUT-8 play central roles in driving recruitment and partitioning within condensates. Despite their importance, the energetic interplay between these residues and their contributions to MUT-16–MUT-8 interactions remain incompletely understood. Computational mutagenesis has emerged as a powerful tool for probing the sequence determinants of phase separation,^36^ while recent applications of double-mutant cycle analysis have enabled the identification of additive versus energetically coupled residue interactions in condensates.^72^ Together, these advances provide a framework for quantitatively dissecting the molecular interactions that govern MUT-16–MUT-8 condensate formation, extending our previous work. ^71^ Going a step further, and given that the material properties of protein condensates play a role in their proper functioning,^16^ this motivates the questioning of how the recruitment of peptides such as the MUT-8 N-terminus changes the material properties of the condensate.

In this study, we systematically investigated the physical properties of phase-separated condensates formed by the MUT-16 M8BR+FFR construct, both in isolation and in the presence of recruited MUT-8 N-terminal peptides. By comparing these systems, we aimed to elucidate how heterotypic interactions influence condensate stability, partitioning behavior, and material characteristics. Firstly, we investigated how MUT-8 recruitment influences the partitioning and conformational properties of MUT-16 across the dilute, dense, and interfacial regions of the condensate. We found that changes in MUT-8 recruitment modulate the conformational behavior of MUT-16 at the interface, as reflected by alterations in the radius of gyration (*R_g_*) and orientational order parameter *S_z_*(*z*). Furthermore, to dissect the molecular determinants underlying recruitment, we employed a double mutant thermodynamic cycle in which targeted mutations were introduced in both the MUT-8 N-terminal sequence and the MUT-16 M8BR+FFR construct.^72^ This framework enabled us to quantify how specific residue-level perturbations affect the transfer free energy, Δ*G*_trans_, of the MUT-16 peptides and determine whether their contributions are additive or energetically coupled. Beyond characterizing recruitment, we further explored whether peptide variants of MUT-8 could modulate the binding affinity of the resident MUT-16 M8BR+FFR chains within the condensate. To this end, we employed an active learning strategy based on Bayesian optimization to identify peptide variants iteratively predicted to render the Δ*G*_trans_ of MUT-16 M8BR+FFR more negative (Fig. 1). Furthermore, our simulations also show the strong correlation previously established by von Bülow *et al.*^35^ between Δ*G*_trans_ and interfacial conformational properties, including *R_g,_*_interface_*/R_g,_*_dense_ and *S_z,_*_interface_. In our case changing the client sequence changes interfacial conformational properties and Δ*G*_trans_ of MUT-16 condensation across all design iterations. Finally, we examined how the compositional evolution of peptide variants across optimization cycles reshapes the material properties of the MUT-16 condensate, thereby linking sequence-level design to emergent condensate behavior. Together, this integrative approach establishes a quantitative framework that connects sequence-level perturbations to interaction energetics and emergent condensate material properties.

**Figure 1:**
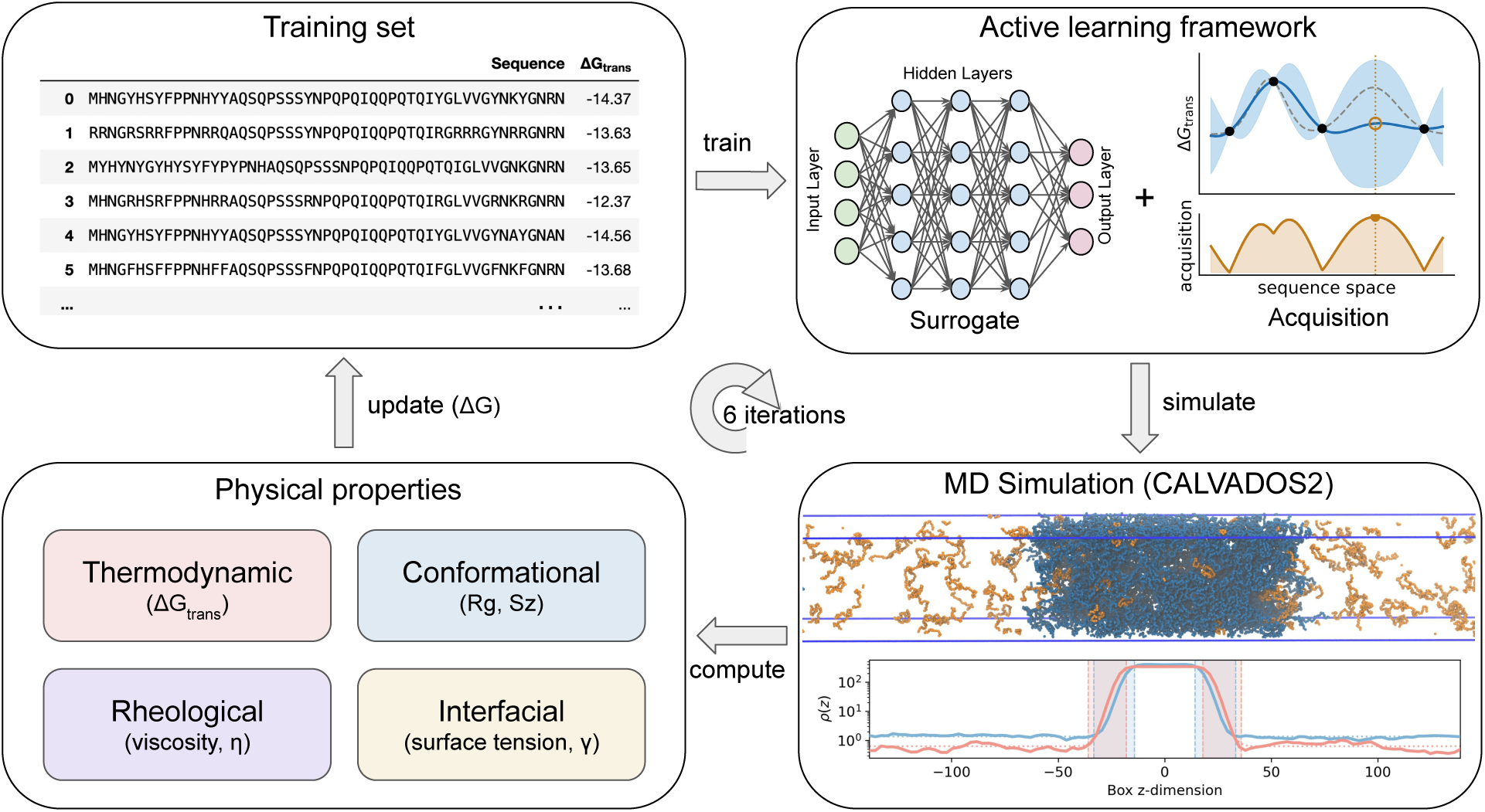
Schematic overview of the active learning loop used in this study to design peptides that modulate the physical properties of the MUT-16 condensate. The loop begins with molecular dynamics (MD) simulations of an initial set of peptide chains coexisting with the condensate. These simulations are used to compute the transfer free energy Δ*G*_trans_, the radius of gyration, the orientation parameter *S_z_*, the viscosity *η*, and the surface tension *γ*. The transfer free energy Δ*G*_trans_, together with its sequence, is added to the training dataset and provided to a Bayesian optimization (BO) framework.^47^ There, sequences are encoded into feature vectors using the UniRep LSTM model,^48^ and a deep ensemble is trained on these features and the computed Δ*G*_trans_ values to learn sequence–property relationships. An acquisition function ranks candidate sequences and proposes new variants expected to improve model performance or target extreme property values. The active learning loop was developed by Yang et al.^47^ and is available through the wazy package. The selected sequences are evaluated via MD simulations, generating training data for the next iteration. This loop is repeated six times, enabling efficient exploration of sequence space while minimising the number of required simulations.

## Methods

### Residue-level coarse-grained simulations

Residue-level coarse-grained simulations^36^ were executed using the CALVADOS2 model^41^ with GPUs using the HOOMD-blue software package (v. 4.5.0).^73^ Within the framework of the CALVADOS2 model, each amino acid is treated as a single bead suspended in an implicit solvent environment. This model enables the investigation of sequence-specific interactions among biomolecules and effectively circumvents the temporal limitations associated with higher-resolution models. The potential energy function employed for the simulations encompasses bonded and non-bonded components, incorporating electrostatic and short-range pairwise interactions. A harmonic potential describes the bonded interactions,

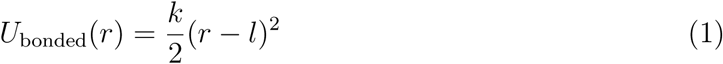

where the spring constant k = 8368 kJ mol^−1^nm^−2^ and equilibrium bond length l = 0.38 nm. The non-bonded interaction between the monomer beads is described by the Ashbaugh-Hatch potential,^74^

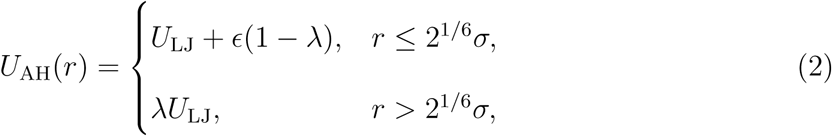

where *ɛ* = 0.8368 kJ mol*−*1 and *U*_LJ_ is the Lennard-Jones potential:

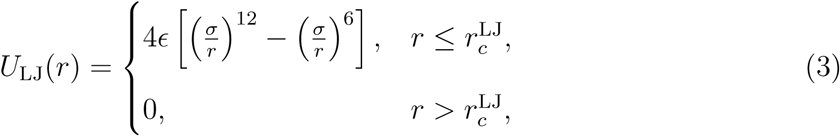

where 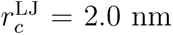.^41^ *σ* and *λ* are determined by computing the arithmetic average of amino acid specific parameters denoting size and hydrophobicity, respectively. The residue-specific *λ* were previously optimized by Tesei et al., with a Bayesian method that uses a comprehensive experimental data set.^41^ The non-bonded interaction also includes an electrostatic component modeled via the Debye-Hückel potential:

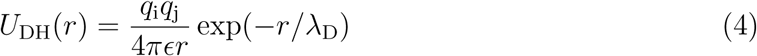

where *q*_i_ and *q*_j_ are charges. The Debye screening length (*λ*_D_) and the dielectric constant (*ɛ*) are set to 1 nm and 80, respectively to reproduce the physiological conditions. The electrostatic potential is truncated at *r_c_* = 4.0 nm.

Unless otherwise specified, the simulations were performed with 100 MUT-16 protein chains in a simulation box of size 20 × 20 × 280nm under periodic boundary conditions, and was run in two replicas. The simulations were initialized by placing the protein chains in the simulation box and running the simulation until a slab forms, for a total of 3 *µ*s. After the MUT-16 slab is formed, 200 peptide chains were added in the simulation box far away from the slab and the system was left to equilibrate. The temperature was maintained at 275 K using a Langevin thermostat. Further, the equations of motions were integrated with a timestep (Δ*t*) of 10 fs. After the system is fully equilibrated, the data is extracted from 1*µ*s runs.

### Density profiles and transfer free energy calculation

To quantify phase separation and peptide partitioning, one-dimensional density profiles were computed along the *z*-axis. Prior to analysis, configurations were recentered such that the condensed phase formed a slab at the center of the simulation box, providing a consistent reference frame for distinguishing dense and dilute regions.

The simulation box was partitioned into bins along *z*, and the local concentration was computed as a function of position to obtain the equilibrium density profile after centering the dense slab in the box. The boundaries between the dense and dilute phases were determined by fitting a hyperbolic tangent to this profile, as described previously,^40,75^

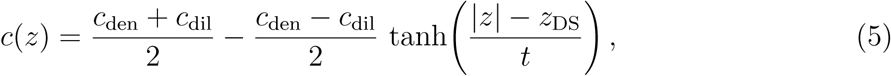

where *z*_DS_ is the position of the dividing surface and *t* the interface thickness. The dense-and dilute-phase regions were taken at cutoffs of |*z*| *< z_DS_* − 1 and |*z*| *> z_DS_* + 4 respectively.

The transfer free energy, Δ*G*_trans_, was calculated as

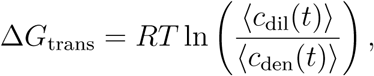

where *R* is the gas constant, *T* is the temperature, and ⟨·⟩ denotes a time average.

### Spatially resolved conformation calculations

To investigate the conformations of the chains in the different phases and at the interface, we calculate the spatially resolved radius of gyration *R_g_*(*z*) and orientational order parameter along the z-direction (normal to the slab interface) *S_z_*(*z*).^35^ We do this for the entire chains, as well as segments of chains. The radius of gyration is calculated as the root-mean-square distance to the center of mass **r**_cm_ for segments composed of *s* residues

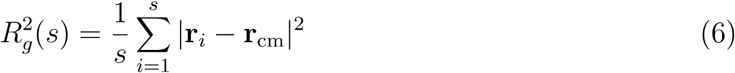

where *s* is the number of residues in a segment and *s* = *N* corresponds to the entire chain, and **r***_i_* is the position of the i-th residue in that segment. The orientational order parameter is calculated through the eigenvectors of the gyration tensor 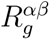

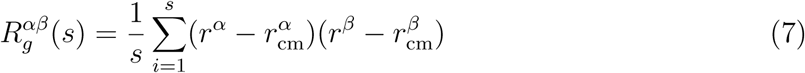

where *α, β* ∈ {*x, y, z*} correspond to the components we are considering. Naturally, we have 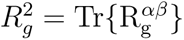. We define the orientation of a segment as the direction of the unit eigenvector **v**^max^ of 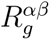 that corresponds to the largest eigenvalue. The orientational order parameter with respect to the z-direction *S_z_* is then calculated as

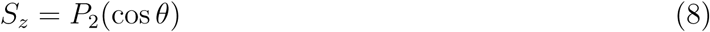

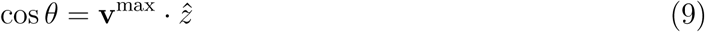

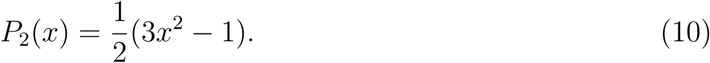

To calculate the Flory exponent *ν*, we exploit the self similarity of the chains and write for the length *R_s_*of segments composed of *s* residues

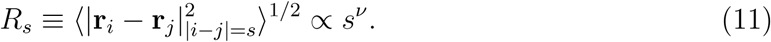

We can then extract the Flory exponent from the average length of *s*-segments by fitting the expected scaling law for different values of *s* ≥ 6.

To obtain a spatial distribution of values for each quantity, we divide our simulation box into equally-sized bins along the z-direction, which is normal to the interface of our condensates. To ensure that our result are robust as to method of assigning values to bins,^76,77^ we try out different methods as detailed below.

The first class of binning relies on the position of the center of mass of the segment.^76^ For each segment composed of *s* residues, we assign the value of *R_g_*(*s*) and *S_z_*(*s*) to the bin that contains its center of mass. To calculate averages, we analyze the trajectory and add values to the corresponding bins while keeping track of the number of segments that were found in that bin, and finally divide by the sample size in each bin. This means that in certain cases few or even no segments are detected in a particular bin, which may result in spurious values or no values at all. This effect is particularly relevant in the bins that correspond to the dilute phases of our systems. With this method, we calculate the profiles using the centers of mass of whole chains (labeled *COM*), as well as segments of the chains where each chain was divided into non-overlapping segments of 26 residues (labeled *L*26) and 52 residues (labeled *L*52).

The second method for binning is to calculate *R_g_* or *S_z_* for each chain, and assign values to the bins based on the occupancy of the beads belonging to that chain, where for each bead we add the value of *R_g_* or *S_z_* to the bin it belongs to.^30,35,77^ At the end of the analysis, the average is calculated as the sample average in each bin. We refer to this as the bead-weighted method and profiles calculated with this methods will be labeled by ‘bead’.

Finally, to assign values to the different phases or to the interface, we fit our density profiles to a hyperbolic tangent as described above. Based on the fit, we divided the system into three regions: an interfacial region defined as |*z*| ∈ [*z_DS_* − *t, z_DS_* + 4*t*], flanked by the dense and dilute phases on either side. We then assign values depending on the phase they belong to.

### Active learning framework for sequence optimization

We consider the problem of mapping a protein sequence to a quantitative property that reflects its interaction with a condensate environment. In this work, the target quantity is the excess transfer free energy, Δ*G*_trans_(MUT − 16), which serves as a measure of MUT-16 chain condensation in the presence of client peptides. The objective is therefore threefold: (i) to learn the relationship between sequence and Δ*G*_trans_, and (ii) to efficiently explore sequence space, and (iii) to identify variants that minimise Δ*G*_trans_, corresponding to enhanced condensation.

To achieve this, we employed the wazy^47^ framework, a Bayesian optimization (BO) approach tailored for protein sequence design. Previously, it was shown that this approach can be meaningfully applied to coarse-grained molecular dynamics simulations of IDPs^26^ (Fig. 1). The workflow is iterative: starting from an initial set of sequences, coarse-grained molecular dynamics simulations are performed to compute Δ*G*_trans_ (MUT-16) in the presence of MUT-8 N-terminal PLD variants. These sequence–property pairs are used to train the wazy^47^ surrogate model, which approximates the mapping from sequence to binding affinity. The trained model is then used to propose new candidate sequences, which are evaluated in subsequent simulation rounds. This loop is repeated over multiple iterations to progressively improve sequence design.

Protein sequences are encoded into fixed-length numerical representations using the UniRep model,^48^ a multiplicative long short-term memory (mLSTM) network trained on a large corpus of protein sequences. This encoding captures sequence-level features relevant for downstream prediction tasks and produces a feature vector of dimension *N* = 1900 for each sequence. These feature vectors, together with the computed Δ*G*_trans_ values, form the input to the wazy^47^ surrogate model.

### Surrogate model and optimization strategy

Within wazy,^47^ the surrogate model is implemented as an ensemble of feedforward neural networks. Each model in the ensemble is a multi-layer perceptron (MLP) that independently predicts the target quantity. In our implementation, the ensemble consists of five MLPs. Each network takes a 1900-dimensional input vector and passes it through three hidden layers with 128, 32, and 2 neurons, respectively. The final layer outputs a predicted mean and standard deviation, representing a Gaussian approximation of the model prediction. Uncertainty estimation is obtained by combining the variance across ensemble predictions with the individual predictive uncertainties.

Sequence selection is guided by an acquisition function that balances exploration of uncertain regions of sequence space with exploitation of sequences predicted to yield favorable properties. In this work, we employ the upper confidence bound (UCB) acquisition function,

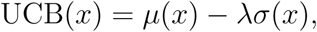

where *µ*(*x*) and *σ*(*x*) denote the predicted mean and uncertainty, respectively, and *λ* controls

the exploration–exploitation trade-off. A value of *λ* = 2 is used throughout.

### Training and validation

The initial training set consists of a combination of manually designed (*ad hoc*) MUT-8 N-terminal PLD mutants and sequences sampled from the ProtGPT protein language model. All sequences are 51 amino acids in length, corresponding to the MUT-8 N-terminal PLD. For each sequence, the corresponding Δ*G*_trans_(MUT − 16) value is computed from coarse-grained molecular dynamics simulations by introducing the mutant peptides into a pre-equilibrated MUT-16 condensate.

To assess model performance, a separate validation set of sequences is constructed using fine-tuned ProtGPT2^59^ (More details in Supporting Information). These sequences are not used during training and serve to monitor predictive accuracy throughout the optimization process. After each optimization step, the neural network model is updated with newly acquired sequence–property pairs, and predictions are evaluated on the validation set using the mean squared error.

Each optimization iteration consists of selecting a batch of candidate sequences^47^ using the acquisition function, evaluating their Δ*G*_trans_ values via simulation, and updating the wazy surrogate model. The optimization objective is to identify MUT-8 variants that induce more negative Δ*G*_trans_ (MUT-16), thereby enhancing the partitioning to the condensate. The initial training phase is referred to as iteration 0, and subsequent iterations progressively refine the sequence–property landscape.

### Viscosity calculation

The viscosity was calculated using equilibrium Green–Kubo relations by integrating the shear stress autocorrelation function *G*(*t*), where we follow the definition of Ref. 78

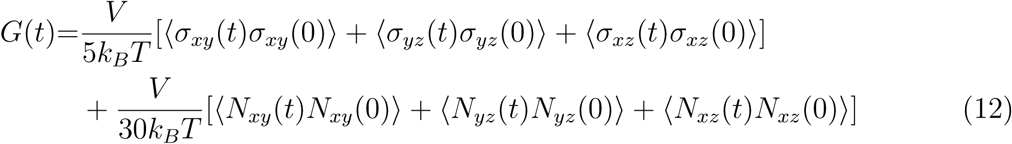

where *σ_αβ_* is the stress tensor, *V* is the volume of the simulation box, *k_B_* is the Boltzmann constant, and *T* is the temperature, and *N_αβ_* = *σ_αα_* − *σ_ββ_*. Simulations were first equilibrated in the NPT ensemble with the barostat set to *P* = 10^−1^ kJ mol^−1^ nm^−3^. Under these conditions, the system evolves into a compact simulation box containing a single condensed phase. At lower pressures, the concentration of peptides in the dilute phase remains significant, which can prevent the formation of a well-defined single-phase system. As our primary objective is to compare the relative effects of different peptide sequences on material properties, the specific choice of pressure should not affect the observed trends. Following equilibration, simulations were performed in the NVT ensemble. The time autocorrelation functions of the stress tensor components were computed at runtime using a multi-tau corre-lator,^78^ implemented to work in HOOMD-blue v4.^79^ The equilibrium viscosity was obtained by integrating the stress relaxation function.^80^ For short times (*t* ≤ 4 × 10^−12^ s), the integral was evaluated numerically using the trapezoidal rule. For longer times (*t* ≥ 4 × 10^−12^ s), the relaxation function was fitted to a sum of Maxwell modes,^80,81^ allowing the remaining contribution to be evaluated analytically. Figure S21A,B shows the stress relaxation function for condensates with and without WT MUT-8 peptides (200 chains). The discrete points correspond to the simulation data, while the squares indicate the fitted Maxwell modes. The dashed line represents the resulting fit used for the analytical evaluation of the long-time contribution to the viscosity.

### Surface tension calculation

The surface tension was calculated from the anisotropy of the stress tensor. Specifically, for a system with two similar planar interfaces oriented normal to the *z*-direction, the surface tension *γ* is given by^82^

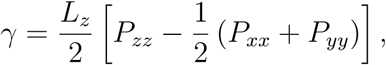

where *P_αα_* are the diagonal components of the pressure tensor and *L_z_* is the box length in the direction perpendicular to the interface. The factor of 1*/*2 accounts for the presence of two interfaces in the simulation box. Simulations were performed in a rectangular box of dimensions 20 × 20 × 280 nm^3^, ensuring sufficient separation between periodic images along the *z*-direction. The stress tensor components were sampled every 0.3 ns, resulting in a total of 10^5^ measurements per simulation, from which time-averaged values of the pressure tensor components were obtained. The statistics of the surface tension measurements are described in the SI.

## Results and Discussion

### Change in the intrinsic properties of a condensate upon addition of MUT-8 peptides

To establish the macroscopic organization of the system, we first computed the density profile of MUT-16 chains along the *z*-axis of the simulation slab, both in the absence and presence of MUT-8 peptides (Fig. 2A). In the pure MUT-16 system, the dense phase exhibited a density of *ρ*_dense_ = 401.11 mg mL^−1^. Following the addition of MUT-8, the dense-phase density of MUT-16 decreased to 339.23 mg mL^−1^ (Fig. 2A), whereas MUT-8 accumulated to a dense-phase concentration of 89.86 mg mL^−1^ (Fig. S1). This shift suggests that incorporation of MUT-8 alters the composition and packing of the condensed phase. A concomitant decrease in MUT-16 density was observed in the dilute phase, where the density dropped from 1.36 mg/ml to 0.63 mg/ml. Despite the simultaneous reduction in both phases, the partition coefficient, defined as *K* = *ρ*_dense_*/ρ*_dilute_, nearly doubled from 294.82 to 536.55 in the presence of MUT-8. This indicates that MUT-8 peptides enhance the relative thermodynamic preference of MUT-16 for the dense phase, even though the absolute density of the condensate is reduced, suggesting that MUT-8 promotes a more strongly partitioned system but loosely packed MUT-16 chains.

**Figure 2:**
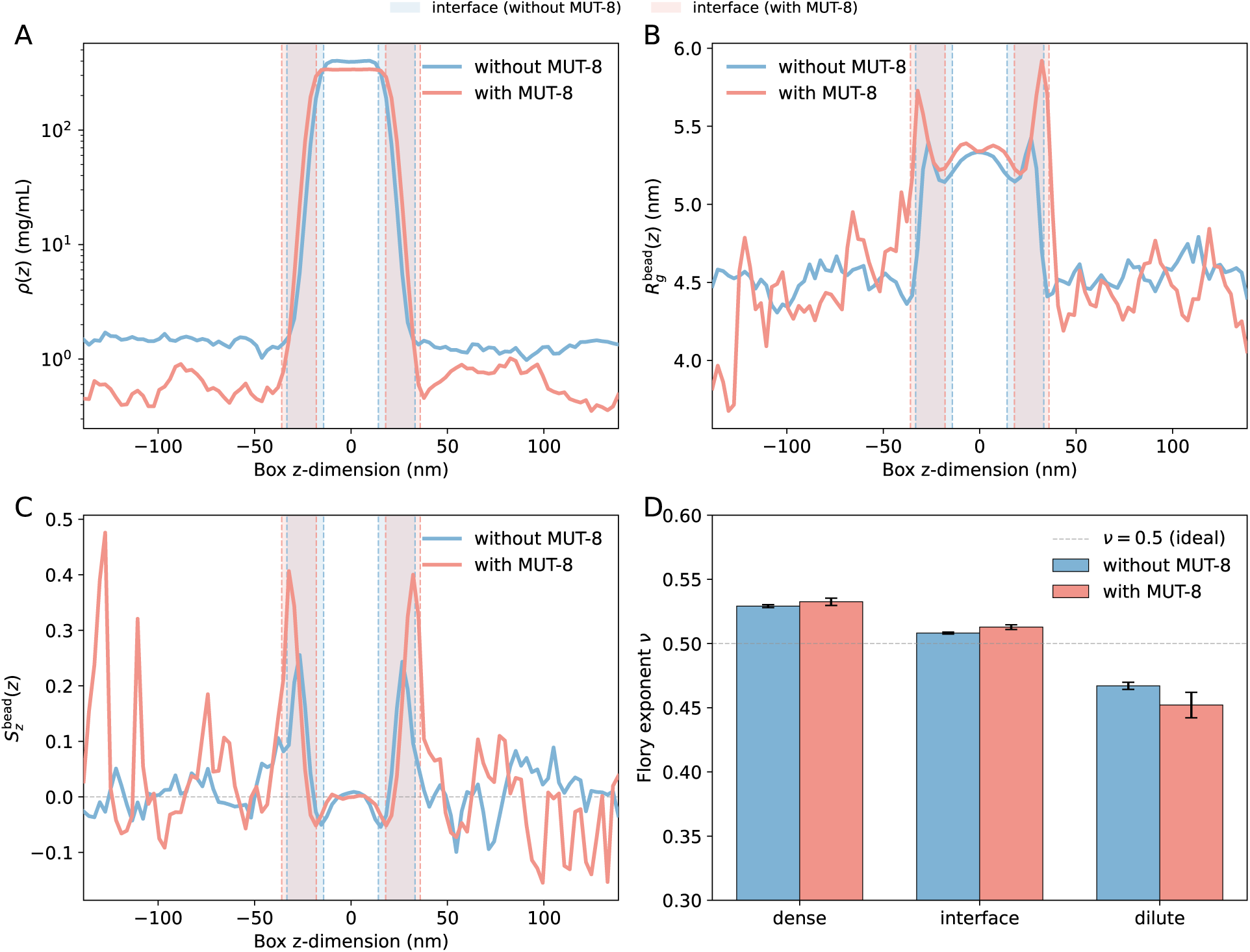
Structural and conformational properties of the MUT-16 condensate with and without MUT-8 peptides. **(A)** Density profile along the *z*-axis of the simulation slab for MUT-16 chains alone (blue) and MUT-16 in the presence of MUT-8 peptides (pink), revealing the dense phase, dilute phase, and interfacial region. Profiles are fitted with a hyperbolic tangent (tanh) function (dashed lines), and the shaded regions between the vertical dashed lines mark the resulting interfaces for each condition. **(B)** Bead-weighted radius of gyration (*R_g_*) profile along the *z*-axis, reporting on the size and compactness of individual chains across the slab. **(C)** Bead-weighted orientational order parameter *S_z_* = ⟨*P*_2_(cos *θ_i_*)⟩ along the *z*-axis, where *θ_i_* is the angle between the principal axis of chain *i* (its longest elongation) and the *z*-axis of the simulation box. *S_z_* quantifies chain alignment along *z*: values near 1 indicate strong alignment with the *z*-axis, 0 corresponds to isotropic orientation, and negative values indicate preferential alignment perpendicular to *z*. **(D)** Flory scaling exponent *ν*, obtained from *R_g_* ∼ *N^ν^*, measured separately in the dense phase, interface, and dilute phase. *ν* reflects the effective solvent quality experienced by the chains, ranging from collapsed (*ν* ≈ 1*/*3) to ideal (*ν* ≈ 1*/*2) to expanded (*ν* ≈ 3*/*5) behavior.

Next we sought to test how the conformations of the scaffold would be affected by the presence of the client. We added 200 MUT-8 peptides (51 aa) to the MUT-16 condensates. We first examined the profile of the radius of gyration *R_g_*(*z*) of MUT-16 chains along the *z*-axis in both conditions. Fig. 2B shows the bead-weighted profile of *R_g_*.^77^ In the absence of MUT-8 peptides, the dense-phase *R_g_* was 5.291 ± 0.004 nm, which increased to 5.347 ± 0.022 nm upon the addition of MUT-8 peptides. Meanwhile, the dilute-phase chains adopted more compact conformations in both systems, with *R_g_* = 4.530 ± 0.018 nm without MUT-8 and 4.434 ± 0.123 nm with MUT-8, indicating that MUT-8 does not appreciably alter the size of isolated MUT-16 chains in the surrounding dilute solution. Using the bead weighted methods, we observe some interesting behavior in the interfacial region and its boundaries as was previously reported.^30^ As we move from inside the condensate into the dilute phase, the *R_g_* has a slight dip both with and without MUT-8, followed by a gradual increase to a peak. However, the peak in *R_g_* in the presence of MUT-8 is significantly larger than without MUT-8 (Fig. 2B). In the bead-weighted profiles, the largest *R_g_* values in both systems were observed in the middle of the interfacial region. The peaks are more pronounced in the presence of MUT-8 (*R_g_* = 5.720 nm at *z* = 32.20 nm) compared to without MUT-8 (*R_g_* = 5.265 nm at *z* = 26.60 nm). Since recent discussion in the literature pointed out that the details of the profile can depend on the choice of binning methods,^30,76,77^ we also considered the profiles obtained through binning based on the center of mass of the chains (Fig. S2A) and for non-overlapping segments of MUT-16 chains composed of 26 residues (Fig. S3A) and 52 residues (Fig. S4A). With these binning methods, the profiles are more noisy due to smaller sample sizes in each bin. In addition, the non-monotonic behavior is much less obvious within our statistical sample. However, regardless of the binning method, MUT-8 chains seem to cause slight swelling of MUT-16. In both of the two replica simulations with see an increase in *R_g_*(*z*) in the interfacial region although it is still not clear whether the peaks in the interface are artifacts due to insufficient sampling.

Next, we analyzed the orientational order parameter relative to the *z*-axis *S_z_*(*z*) of MUT-16 chains to characterize how the chains are aligned relative to the slab normal in both conditions. Fig. 2C shows the bead-weighted profiles for *S_z_*. In the dense phase, *S_z_* was essentially zero in both systems (−0.0002±0.0086 without MUT-8 and −0.0027±0.0053 with MUT-8), indicating that MUT-16 chains adopt isotropic orientations within the condensate interior regardless of the presence of MUT-8, in correspondence to previous results.^29^ As we move from the dense into the dilute phase, *S_z_* shows non-monotonic behavior. At the inner boundary of the interface, *S_z_* takes on slightly negative values, signaling a slight tangential alignment of the chains.^29^ This is followed by gradual increase into positively valued peaks in *S_z_* inside the interfacial region. This indicates that the chains are preferentially aligning perpendicular to the interface of the condensate. While the slight tangential alignment is not affected by the presence of MUT-8, the perpendicular alignment increases roughly by a factor of two when MUT-8 peptides are introduced. In the dilute phase, despite large fluctuations due to poor statistics, we recover isotropic alignment of the chains. To see the effect of the binning method on the orientation profile, we also calculated the profiles using the center of mass binning method (Fig. S2B) and for 26-residues segments (Fig. S3B) and 52-residue segments (Fig. S4B). The profiles are again more noisy, yet show similar features to the bead-weighted profiles, especially as we transition from the dense phase into the interface, with both replica simulations showing similar behavior. These profiles show a drop of *S_z_*(*z*) at the inner boundary of the condensate, with *S_z_*(*z*) turning positive at the interface.

Finally, to interpret these conformational changes in terms of the effective solvent quality experienced by MUT-16 chains, we computed an effective Flory scaling exponent, *ν*, from the scaling relation *R_s_* ∼ *s^ν^* (see Methods section) separately within the dense phase, the interface, and the dilute phase where values were assigned based on the position of the centers of mass of the chains(Fig. 2D). The value of *ν* provides a direct readout of the effective solvent quality: *ν* ≈ 0.333 corresponds to collapsed, globule-like chains in a poor solvent, *ν* = 0.500 to ideal-chain (theta-solvent) behavior, and *ν* ≈ 0.588 to expanded, self-avoiding chains in a good solvent.^83^ In the dense phase, *ν* was 0.5291 ± 0.0011 without MUT-8 and increased slightly to 0.5325 ± 0.0028 in the presence of MUT-8 but stayed withing the confidence interval, indicating that MUT-16 chains experience near-ideal, theta-like conditions within the condensate both with and without MUT-8. At the interface, *ν* remained near-ideal in both conditions, with *ν* = 0.5082 ± 0.0007 without MUT-8 and *ν* = 0.5128 ± 0.0018 with MUT-8 but there is a small statistically significant increase upon addition of MUT-8. In contrast, the dilute-phase *ν* decreased from 0.4671 ± 0.0028 without MUT-8 to 0.452 ± 0.010 with MUT-8, moving further away from the ideal value and toward more compact conformations.

In conclusion, these analyses reveal that MUT-8 peptides can reshape the conformational landscape of MUT-16 in a phase-dependent manner. Within the dense phase, MUT-8 lowers the bulk concentration of MUT-16 while allowing MUT-16 chains to adopt slightly more expanded conformations, indicating that MUT-8 modulates MUT-16 packing without disrupting the condensed state. At the outer boundary of the interface, all approaches indicate some preferential alignment in the direction perpendicular to the interface for MUT-16, while chains are orientated tangentially to the interface at the inner boundary of the interface, in agreement with previous results.^29,30,35^ The perpendicular alignment seems to be amplified by MUT-8 and maximized closer to the outer edge. In the dilute phase we found that MUT-8 slightly lowers the Flory exponent. Compared with the behavior of the Flory exponent in the dense phase upon addition of MUT-8, this opposite effect on the effective solvent quality across the two phases provides an explanation for the doubling of the partition coefficient. Making the dilute phase a poorer solvent environment directly amplifies the thermodynamic preference of MUT-16 for the condensate. Collectively, these results indicate that MUT-8 not only redistributes MUT-16 between the two phases but also locally restructures the condensate by promoting slightly more expanded, oriented MUT-16 chains at the interface, with potential consequences for the mechanical and interfacial properties of the condensate.

### Effect of client mutations on scaffold partitioning

To chemically dissect the mechanism underlying the recruitment of MUT-8 into MUT-16 condensates, we focused on the specific intermolecular interactions that mediate this process. Previous coarse-grained and atomistic simulations, together with *in vitro* experiments,^71^ identified interactions between Arg residues in the MUT-16 M8BR region and Tyr residues in the N-terminal domain of MUT-8 as key determinants of recruitment. Experimentally, progressive mutation of Arg residues in MUT-16 M8BR to Lys or Ala resulted in a systematic reduction in MUT-8 binding, indicating the importance of these residues.^71^ Complementary simulation analysis revealed that Arg engages Tyr through a combination of cation–*π*, *π*–*π*, and hydrogen-bonding interactions involving the Tyr backbone.^10,13,71^ The multivalent, tridentate character of this interaction motif provides a strong and structurally favorable binding interface, thereby promoting efficient recruitment of MUT-8 into the MUT-16 condensate.

To further probe whether mutations in the client (MUT-8) sequence can modulate phase separation of the scaffold (MUT-16), we systematically introduced a set of targeted mutations designed to test how specific residue changes in the client affect MUT-16 partitioning, quantified by the transfer free energy Δ*G*_trans_(MUT-16).

The *ad hoc* mutations were first designed to further probe the role of Tyr residues in MUT-8 (WT) by substituting them with Phe (Y2F), Ala (Y2A), Asp (Y2D), Glu (Y2E), Arg (Y2R), and Lys (Y2K). We further investigated the spatial distribution of Tyr residues by redistributing them in distinct patterns: evenly spaced along the sequence (Y-spaced), clustered at the N-terminus (Y-N cluster), or clustered at the C-terminus (Y-C cluster). To examine the role of charge, we generated a neutral variant by mutating all charged residues to Ala. In addition, we constructed fully positively and negatively charged variants by mutating residues to Arg and Glu, respectively, while preserving Gln, Asn, Gly, Ser, and Pro (Fig. 3A). These residues were retained due to their known importance in prion-like domain (PLD) low-complexity regions.^13^ Peptide chains carrying the above-mentioned mutations were introduced into a pre-equilibrated MUT-16 condensate (Methods), and the resulting change in the excess transfer free energy, Δ*G*_trans_(MUT-16) was subsequently calculated.

**Figure 3:**
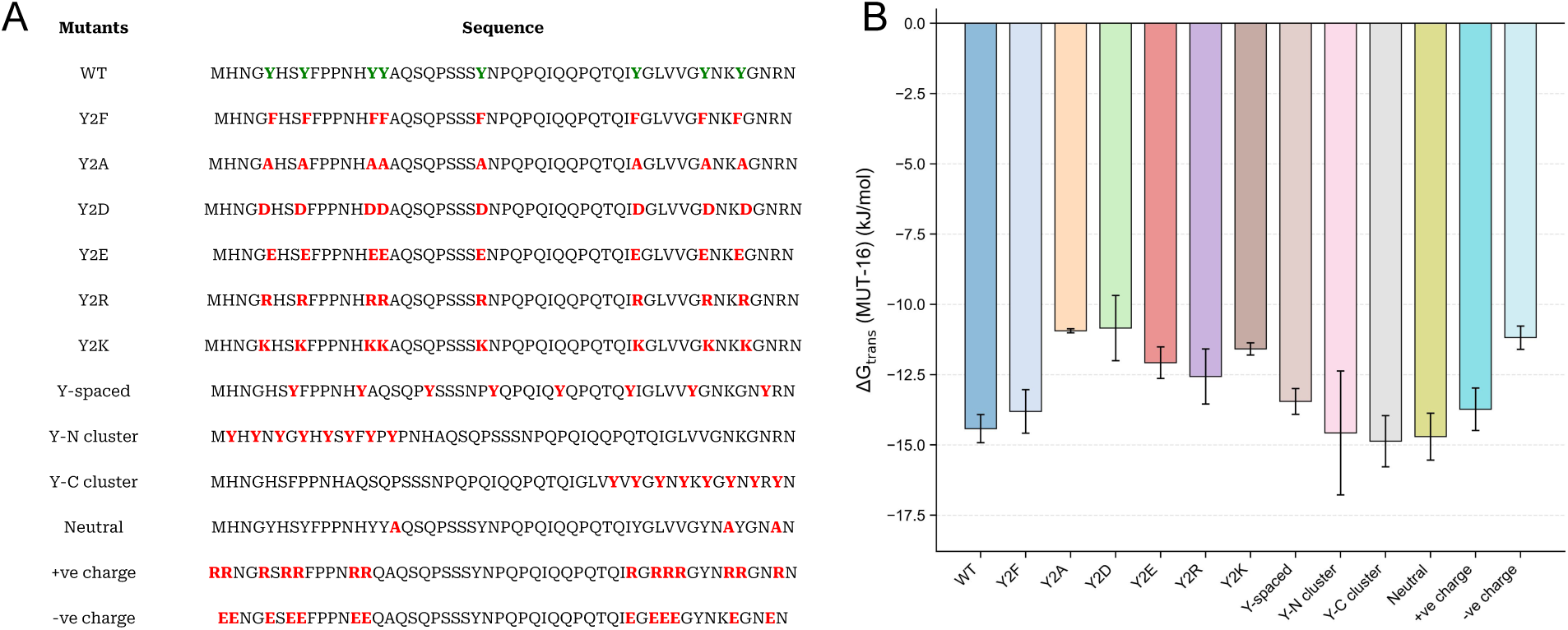
The N-terminal PLD of MUT-8 WT and a series of designed *ad hoc* variants were investigated to assess their effect on MUT-16 condensate recruitment. For each sequence, the phase separation propensity of MUT-16 was quantified by the transfer free energy of MUT-16 chains upon recruitment, Δ*G*_trans_(MUT-16). **A.** To probe the role of the tyrosine residues critical for recruitment, all Tyr were substituted with Phe (Y2F), Ala (Y2A), Asp (Y2D), Glu (Y2E), Arg (Y2R), or Lys (Y2K). Patterning effects were examined by redistributing Tyr uniformly (Y-spaced) or clustering them at the N- (Y-N cluster) or C-terminus (Y-C cluster). Charge effects were probed with variants of altered net charge: a neutral variant (Neutral) replacing charged residues with Ala, a positive variant (+ve charge) using Arg, and a negative variant (-ve charge) using Glu; in all three, Gln, Asn, Gly, Ser, and Pro were preserved to retain the intrinsic properties of the low-complexity PLD. **B.** Bar plot of Δ*G*_trans_(MUT-16) upon recruitment of MUT-8 WT and each variant.

We observed that the Y2F mutation has a negligible effect on the transfer free energy,= Δ*G*_trans_ (MUT-16), remaining within error of the WT. In contrast, mutations to Ala (Y2A) and Asp (Y2D) significantly reduce Δ*G*_trans_ (MUT-16) (Fig. 3B). The Y2R mutant did not show a significant change compared to WT, while Y2K and Y2E showed a moderate statistically significant reduction. For Tyr redistribution, none of the rearrangements resulted in statistically significant changes in Δ*G*_trans_ from its value for WT. For charge-modified variants, the neutral peptide and positively charged variants show behavior comparable to WT, whereas the negatively charged variant yields a less negative Δ*G*_trans_, indicating reduced binding affinity compared to WT (Fig. 3B).

Motivated by the findings of Rauh *et al.*,^72^ we tested whether scaffold (MUT-16) and client (MUT-8) mutations exert additive effects on phase behavior by comparing double mutants with the corresponding single mutants (Fig. S5; Fig. S6). If scaffold and client mutations act independently, the shift in Δ*G*_trans_ for the double mutant should equal the sum of the shifts measured for each single mutant. A deviation from this additive prediction would instead indicate coupling between the two mutations. Across all systems examined, the effects were largely additive, with ΔΔ*G*_int_ ≈ 0, indicating little to no energetic coupling between mutations on the scaffold and the client, even for mutations involving residues that participate in cation–*π* interactions.

In conclusion, mutations in the client (MUT-8), particularly those targeting aromatic residues, can modulate the transfer free energy, Δ*G*_trans_, of the scaffold (MUT-16). Together with the observed additivity of scaffold and client mutations, these results suggest that condensate partitioning can be tuned predictably through sequence modifications, with individual perturbations contributing approximately independently.

### Design of peptide variants using active learning to modulate MUT-16 partitioning

Building on these insights, we employed the active-learning framework wazy,^47^ which combines a pre-trained protein language model with Bayesian optimization to iteratively propose and evaluate new peptide variants of the same length as MUT-8. With this approach we want to find peptides that enhance MUT-16 condensation. In this approach, each candidate sequence is first converted into a fixed-length numeric vector using the pre-trained UniRep model,^48^ which is then used to train a neural network ensemble that learns how sequence relates to Δ*G*_trans_(MUT-16). The model then proposes the next set of sequences to simulate, focusing on the most informative candidates so that sequence space can be explored efficiently from a limited number of labeled examples. Iterating this cycle identified variants predicted to enhance MUT-16 partitioning into the condensate, reflected by more negative values of Δ*G*_trans_(MUT-16). Across successive wazy^47^ iterations, we simulated the MUT-16 condensate in the presence of each proposed peptide variant (20 per round), with each simulation containing 100 MUT-16 chains and 200 variant chains, and computed the corresponding Δ*G*_trans_(MUT-16) values (Fig. 1).

Over the course of training, the surrogate model became progressively better at predicting Δ*G*_trans_(MUT-16): the Pearson correlation coefficient between predicted and measured values rose steadily from *r* = 0.22 at iteration 0 to a peak of *r* = 0.84 at iteration 6 (Fig. S8). Consistent with this improving fit, the prediction error on Δ*G*_trans_(MUT-16) as modeled by the ensemble of neural networks decreased across iterations.^47^ To assess generalization beyond the iteratively sampled sequences, we additionally overlaid a test set of 20 sequences generated by the fine-tuned ProtGPT2 model (Fig. S7). We first observed that the active-learning set comprised stronger interactors than the independent set, with more negative Δ*G*_trans_(MUT-16) values: the interquartile range spanned −14 to −17 kJ/mol for the active-learning set, compared with −12 to −14 kJ/mol for the independent set. Furthermore, the surrogate model’s predictions for the independent set also improved over successive iterations, indicating that the learned sequence–property relationship transfers to sequences drawn from a different generative source.

The changes in Δ*G*_trans_ over the course of the active learning are summarized as box- and-whisker plots with overlaid individual data points (Fig. 4A). In the initial selection, the distribution was relatively broad, with an interquartile range (IQR) spanning roughly −11 to −13.5 kJ/mol and a median near −12.2 kJ/mol. The median remained comparable through iteration 3, fluctuating around −12 kJ/mol, before shifting markedly downward from iteration 4 onward to approximately −16 to −18 kJ/mol, indicating a substantial enhancement of MUT-16 partitioning. The spread of the distribution evolved in parallel: the IQR narrowed in iteration 1 and remained narrow through iteration 3, consistent with convergence of the search toward a localized region of sequence space, then widened in iterations 4 and 5, reflecting renewed exploration, before narrowing again in iteration 6. Iteration 6 was also the only round in which a substantial fraction of simulations failed to converge (10 of 20), owing to slow relaxation of client clusters. To assess whether this reflected an intrinsic change in the variants, we estimated their phase-separation propensity using the predictor of von Bülow *et al.*,^40^ computing the predicted Δ*G*_trans_ and saturation concentration for each variant. By iteration 6, the variants had acquired the capacity for self-condensation: the predicted Δ*G* exceeded 2 *k*_B_*T* and the saturation concentration approached zero (Fig. S9A,B). This emergence of intrinsic phase separation provided a natural stopping point, and we terminated the optimization after iteration 6.

**Figure 4:**
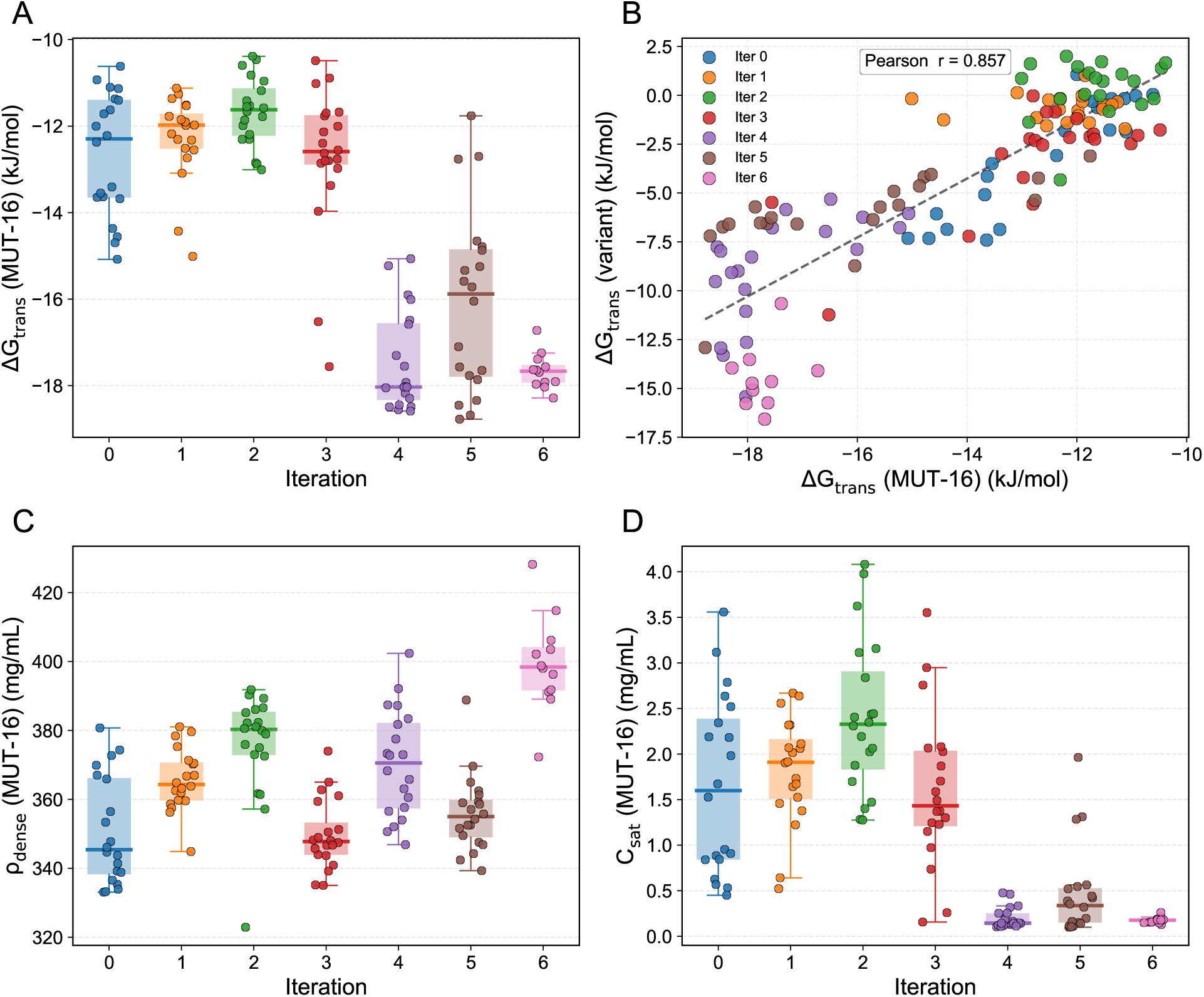
Active-learning optimization of peptides (variants) that promote MUT-16 condensation upon recruitment. Using the wazy^47^ Bayesian-optimization framework, new peptide variants of the same length as MUT-8 WT were proposed at each iteration, simulated, and used to retrain the model. The objective was to identify peptides whose recruitment drives the scaffold transfer free energy, Δ*G*_trans_(MUT-16), toward more negative values, reflecting enhanced MUT-16 phase separation. In panels **A**, **C**, and **D**, the distribution of each metric across successive iterations is shown as a box-and-whisker plot (central line, median; box, interquartile range (IQR); whiskers, data within 1.5×IQR), with individual measurements overlaid as scatter points. **A.** Δ*G*_trans_(MUT-16). **B.** Correlation between Δ*G*_trans_(MUT-16) and the variant transfer free energy Δ*G*_trans_(variant). **C.** Dense-phase density, *ρ*_dense_(MUT-16). **D.** Saturation concentration, *C*_sat_(MUT-16).

Δ*G*_trans_(MUT-16) and Δ*G*_trans_(variant) were strongly correlated (Pearson *r* = 0.857; Fig. 4B): client variants that partition more favorably into the condensate (Fig. S10) are also those in which the scaffold condenses most strongly. This is consistent with both quantities being governed by the same scaffold–client binding affinity. The dense-phase density *ρ*_dense_(MUT-16) (Fig. 4C) showed no systematic trend, with the median fluctuating from iteration to iteration. In contrast, the median saturation concentration (*C*_sat_(MUT-16)) decreased steadily across iterations (Fig. 4D), which renders Δ*G*_trans_(MUT-16) more favourable. Building on this picture, we examined how the interfacial conformational properties of MUT-16 evolve across the peptide-design iterations. We tracked the ratio of the interfacial to dense-phase radius of gyration, *R_g,_*_interface_*/R_g,_*_dense_ (Fig. S11A), and the interfacial orientational order parameter, *S_z,_*_interface_ (Fig. S11B), taking the peak interfacial values from the bead-weighted profiles in each case.^35^ Over the iterations, with median *R_g,_*_interface_*/R_g,_*_dense_ increasing from near 1 toward ∼ 1.35 (Fig. S11A), indicating that MUT-16 chains progressively expand at the interface relative to their more compact conformations in the dense-phase interior. In parallel, median *S_z,_*_interface_ rose from near 0.2 toward ∼ 0.75 (Fig. S11B) over the iterations, signifying a shift from essentially isotropic chain orientations to strong alignment along the slab normal at the interface. Similar behavior can be observed for the conformations obtained using other binning methods, as can be seen in Figs. S12-S14. There, we again defined the interfacial values as those corresponding to peaks in the interface, and we ensured that the detected peak is not the result of a single measurement. The profiles for all the mutants using all the binning methods we tried, along with the detected peaks can be found within the data repository associated with this article.

We then computed the correlation of these two descriptors with Δ*G*_trans_(MUT-16) across the peptides from active learning cycles. For the values obtained from bead-weighted profiles, both descriptors showed strong negative correlations with Δ*G*_trans_(MUT-16): *R_g,_*_interface_*/R_g,_*_dense_ (Pearson *r* = −0.900; Fig. 5A) and *S_z,_*_interface_ (*r* = −0.919; Fig. 5B). Thus, peptide variants that drive MUT-16 chains to expand and orient perpendicular to the interface also yield a more negative transfer free energy, and therefore stronger partitioning into the condensate as previously established by von Bülow et al.^35^ The two descriptors were themselves highly correlated (*r* = 0.970; Fig. 5C), indicating that interfacial chain expansion and alignment are tightly coupled across the ensemble. Together, these results suggest that the iterative design progressively reshapes MUT-16 conformations at the interface in step with its strengthening thermodynamic preference for the dense phase. The correlations between the conformations and Δ*G*_trans_(MUT-16) remain present regardless of the binning method we used although they are less pronounced (Figs S15-S17).

**Figure 5:**
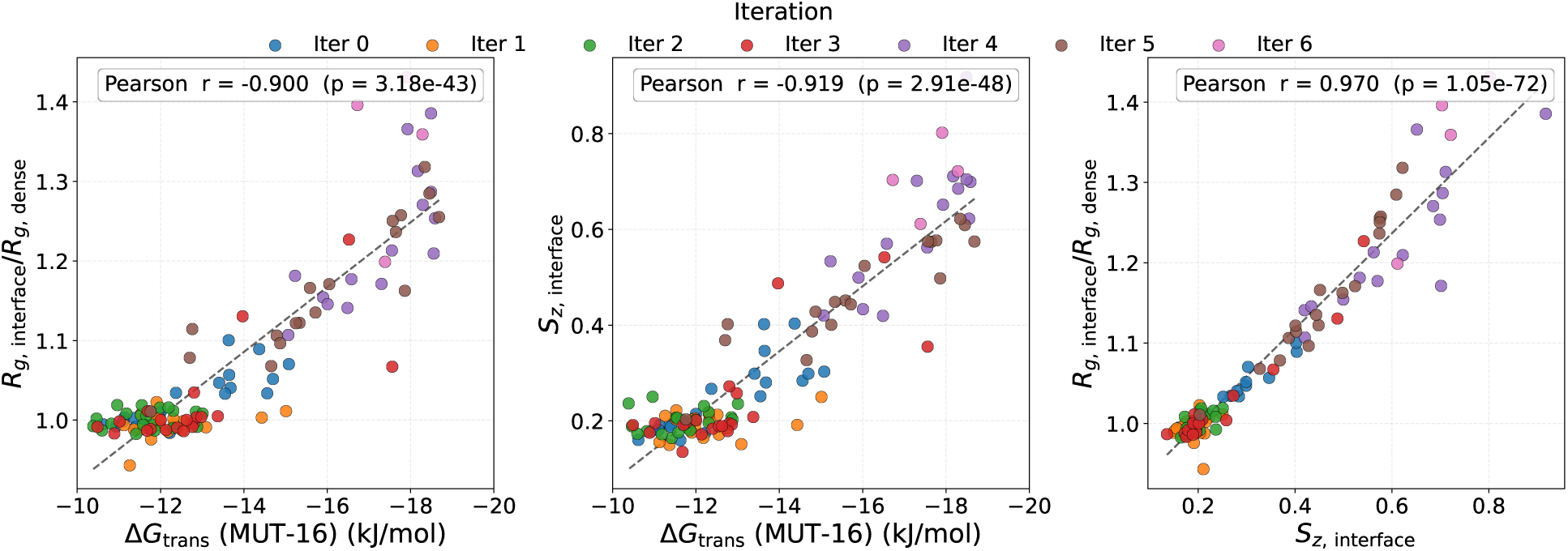
Correlations between the interfacial conformational properties of MUT-16 chains from bead-weighted profiles and the transfer free energy Δ*G*_trans_(MUT-16), evaluated across peptide variants from successive design iterations. The ratio *R_g,_*_interface_*/R_g,_*_dense_ reports how much MUT-16 chains expand or contract at the interface relative to the condensed interior, *S_z,_*_interface_ is the orientational order parameter of interfacial chains, and Δ*G*_trans_(MUT-16) quantifies the preference of MUT-16 for the dense over the dilute phase. **(A)** *R_g,_*_interface_*/R_g,_*_dense_ vs. Δ*G*_trans_(MUT-16). **(B)** *S_z,_*_interface_ vs. Δ*G*_trans_(MUT-16), relating interfacial chain alignment to the transfer free energy. **(C)** *R_g,_*_interface_*/R_g,_*_dense_ vs. *S_z,_*_interface_, relating chain expansion to orientational ordering at the interface. Each point is a distinct peptide variant from the iterative design procedure; the Pearson correlation coefficient is shown in each panel.

We next quantified how the residue-class composition of the proposed peptides shifts across iterations, grouping amino acids into hydrophobic, polar uncharged, charged, and aromatic categories (Fig. S18). The initial sequences (iteration 0) were predominantly hydrophobic and polar, with median counts of ∼20 hydrophobic, ∼20 polar uncharged, ∼10 charged, and only ∼1 aromatic residue. Through iterations 1 to 3, the hydrophobic and polar uncharged counts fluctuated without a sustained trend, charged residues dipped and recovered, and aromatic content rose only modestly (Fig. S18A–D). The decisive changes emerged in iterations 4 to 6: hydrophobic and polar uncharged residues were strongly depleted, with both medians and IQRs falling below five (Fig. S18A,B); charged residues peaked near a median of ∼13 at iteration 4 before settling to ∼10 (Fig. S18C); and aromatic residues were dramatically enriched, with the median rising to ∼17 and the upper quartile reaching ∼22 (Fig. S18D). These trends point to a selection pressure favoring aromatic residues and a moderate charged content while depleting generic hydrophobic and polar residues, consistent with the increasingly negative Δ*G*_trans_(MUT-16) (Fig. 4A) and Δ*G*_trans_ of the peptide variants (Fig. S10) reached over the same iterations.

Given this enrichment of charged residues, we further resolved the charge composition into net charge per residue (NCPR), the fraction of charged residues (FCR), and its positive (FPCR) and negative (FNCR) components (Fig. S19). Notably, MUT-16 M8BR+FFR is overall charge-balanced, with an NCPR of 0.00 and an FCR of 0.12. NCPR of the peptide variants began near a median of ∼0.1 with a broad IQR and varied non-monotonically across iterations, decreasing to iteration 2, rising through iteration 4, then declining slightly by iteration 6, indicating dynamic tuning of net charge rather than a steady trend (Fig. S19A). The total FCR showed no consistent direction (Fig. S19B), but its decomposition was more revealing: FPCR rose to a peak at iteration 4 before easing in the final rounds, likely a compositional trade-off against the strong aromatic enrichment in iterations 5-6 (Fig. S19C), whereas FNCR fell progressively to near zero (Fig. S19D). We further examined the increase in positive charge count across iterations (Fig. S20A) and found it to be driven by arginine (Fig. S20B) rather than lysine (Fig. S20C). This preferential enrichment of arginine over lysine, despite their identical formal charge, suggests that arginine is selected primarily as a sticker rather than merely as a positive charge.

In conclusion, the active-learning framework efficiently designed peptide clients that progressively strengthen the phase separation of the MUT-16 scaffold upon recruitment. Over the iterations, Δ*G*_trans_(MUT-16) became increasingly negative, with a rising partition coefficient and a falling saturation concentration. These thermodynamic gains correlated with the swelling and alignment of MUT-16 chains at the condensate interface. At the sequence level, the designed clients showed a compositional shift toward aromatic residues and positive charge, at the expense of negative, hydrophobic, and polar residues, which is consistent with the known sequence determinants of phase separation.^10,40^ Together, these results establish active learning as a practical route to design condensate-modulating peptides while recovering the sequence determinants of phase behavior.

### Peptide variants modulate the material properties of the condensate upon recruitment

To examine how recruited peptide variants modulate the material properties of the MUT-16 condensate, we computed the viscosity and surface tension for three representative variants from each iteration of the active-learning framework. These variants were selected based on exhibiting the most negative Δ*G*_trans_ (MUT-16) upon recruitment into the condensate. Both properties were derived from the components of the stress tensor, which were evaluated at runtime using built-in methods in HOOMD-blue.^73^ The surface tension was obtained from the anisotropy of the stress tensor. For this purpose, the stress tensor was sampled every 0.3 ns, yielding a total of 10^5^ measurements per simulation (see Supporting Information for details on statistics). The viscosity was calculated using equilibrium Green–Kubo relations by integrating the shear stress autocorrelation function, also known as the shear stress relaxation function, *G*(*t*). To calculate *G*(*t*) we employ a multi-tau correlator^79^ to calculate correlations on the fly in the simulation, including measurements from every time step. For each system, two independent simulations were performed to ensure statistical reliability. Each simulation contained 100 MUT-16 chains and 50 chains of peptide variants. All simulations were run for 3 *µ*s, starting from configurations that had been pre-equilibrated for at least 1 *µ*s.

We observed that the viscosity of the MUT-16 condensate is markedly reduced upon the addition of WT MUT-8 N-terminal PLD peptide chains. Specifically, the viscosity decreases from *η* = 4.60 × 10^−6^ Pa · s in the absence of peptides to *η* = 2.33 × 10^−6^ Pa · s upon their inclusion (Fig. 6A), highlighting the pronounced impact of client recruitment on condensate material properties. Across the wazy^47^ iterations, the average viscosity calculated from the three selected MUT-8 mutants exhibits an overall increasing trend until iteration 4, then stabilizes in later iterations (Fig. 6A). To quantify the relationship between client mutations and condensate mechanics, we evaluated the correlation between Δ*G*_trans_ (MUT-16) and condensate viscosity across iterations (Fig. 6C). We find a strong negative correlation, with a Pearson correlation coefficient of *r* = −0.813, indicating that mutations promoting stronger interactions (more negative Δ*G*_trans_(*MUT* − 16)) are associated with more viscous condensates.

**Figure 6:**
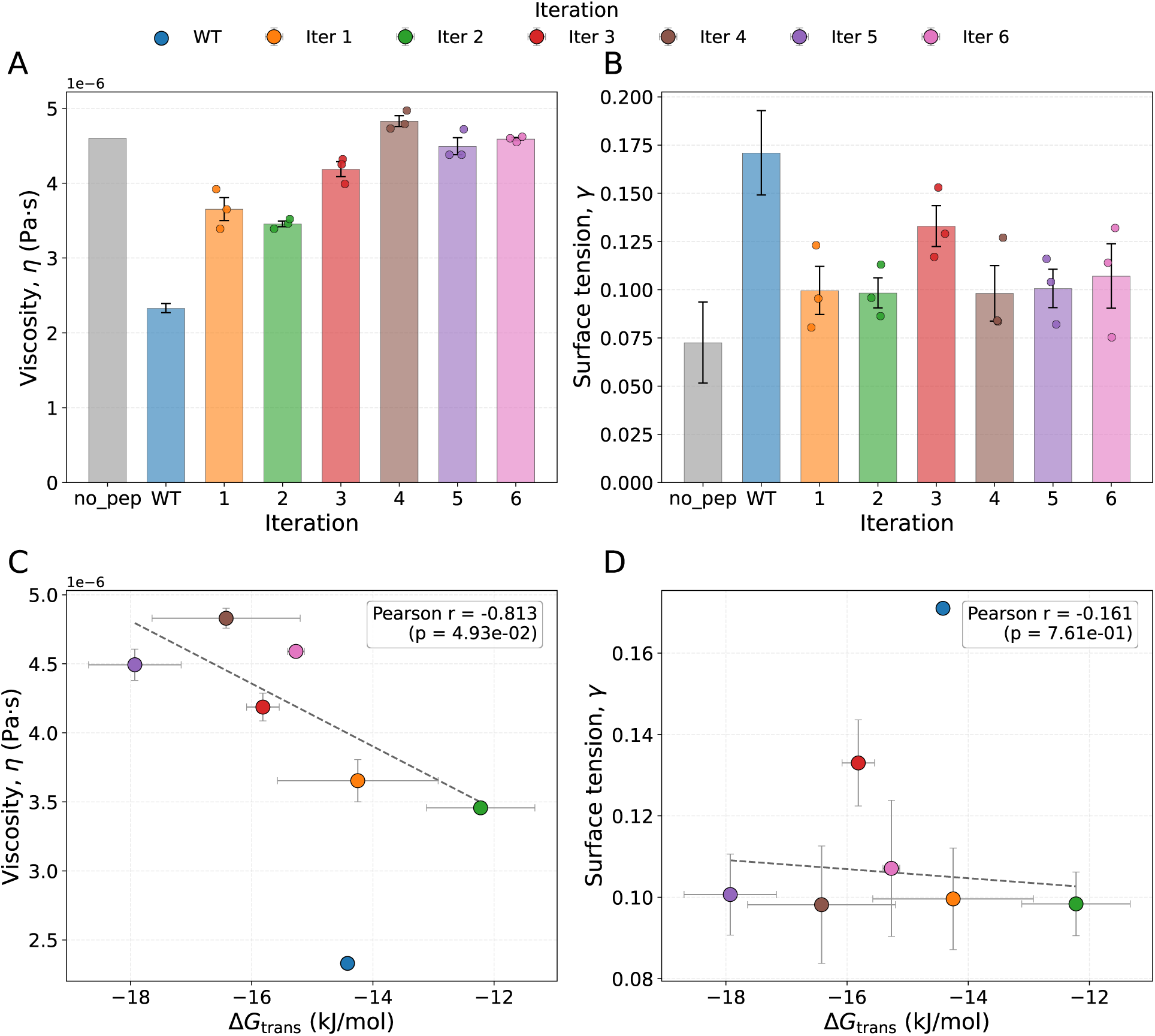
Material properties of the MUT-16 condensate without peptide (no_pep), with WT MUT-8, and for three representative peptide variants per wazy^47^ training iteration, selected as the sequences with the most negative Δ*G*_trans_(MUT-16). For each iteration, the bar shows the mean with error bars, and individual values are overlaid as scatter points. **A.** Viscosity. **B.** Surface tension. **C.** Correlation between Δ*G*_trans_(MUT-16) and viscosity. **D.** Correlation between Δ*G*_trans_(MUT-16) and surface tension. Dashed black lines in (C) and (D) show the linear regression.

In contrast to viscosity, the surface tension of the MUT-16 condensate increases markedly upon the addition of WT MUT-8 peptides, rising from *γ* = 7.25 × 10^−2^ mN*/*m to *γ* = 17.1 × 10^−2^ mN*/*m. Across the wazy^47^ iterations, the average surface tension of the three selected mutants shows an unchanging trend, as the majority of the values fall within each other’s confidence intervals. Correlation analysis reveals no relationship between Δ*G*_trans_ (MUT-16) and surface tension (Fig. 6D), with a Pearson coefficient of *r* = −0.161.

Given the strong correlations of both the interfacial *R_g_* ratio, (*R_g,_*_interface_*/R_g,_*_dense_), and *S_z,_*_interface_ with Δ*G*_trans_ (MUT-16) established above, we next asked whether these interfacial conformational descriptors are also predictive of the material properties of the MUT-16 condensate. To this end, we calculated these quantities for the condensates in the presence of 50 peptides and examined their correlations with the condensate viscosity (*η*) and surface tension (*γ*) (Fig. S24). The change in interfacial conformations across different iterations shows similar trends for 50 peptides as it did for 200 peptides. In addition, both descriptors are positively correlated with viscosity, with Pearson *r* = 0.764 for the *R_g_* ratio (Fig. S24A) and Pearson *r* = 0.887 for *S_z,_*_interface_ (Fig. S24B), indicating that peptide variants which promote interfacial stretching and alignment of MUT-16 chains also yield more viscous condensates. In contrast, neither descriptor showed an appreciable correlation with surface tension, with Pearson *r* = 0.072 for the *R_g_* ratio (Fig. S24C) and Pearson *r* = 0.226 for *S_z,_*_interface_ (Fig. S24D). This is particularly interesting since the peptides affect the interfacial conformations of the chains, yet those conformational changes do not lead to a change in the interfacial tension, or their effect is compensated for by other mechanisms. The exception to this trend, is the increase in surface tension upon addition of WT MUT-8 peptides, which also changes the interfacial conformation of the chains (Fig. 2, Fig. S2, Fig. S3, Fig. S3).

In conclusion, the recruitment of MUT-8 WT into the condensate markedly altered the viscosity of MUT-16 condensates, but did not affect the surface tension except in the case of WT MUT-8 peptides. Across the iterations, the viscosity increased gradually, whereas the surface tension showed no clear trend. The viscosity correlated strongly with both the scaffold partitioning and the interfacial conformations of MUT-16 chains, while the surface tension was essentially uncorrelated with either.

## Conclusions

In this work, we show that the physical properties of a phase-separated protein condensate can be tuned by introducing peptide clients engineered to bind the scaffold that constitutes the condensate. Using this strategy, we highlight how a designed client reshapes scaffold conformations, partitioning, and bulk material behavior, and we establish a pipeline for generating such modulators.

Our analyses show that MUT-8 WT peptides remodel the MUT-16 scaffold in a phase-dependent manner. In the dense phase, MUT-8 lowers the bulk concentration of MUT-16 and allows slightly more expanded chains, loosening the packing of the scaffold in the condensate without dissolving it. Meanwhile, at the interface, the addition of MUT-8 promotes weak alignment along the direction normal to the slab. Finally, in the dilute phase, *R_g_* is essentially unchanged except for a slight decrease in the Flory exponent *ν*, signaling an effectively poorer solvent environment. Because the effective solvent quality worsens in the dilute phase yet at worst remains unchanged in the dense phase, the thermodynamic preference of MUT-16 for the condensate increases, explaining the roughly twofold rise in the partition coefficient upon addition of MUT-8. Beyond redistributing scaffold between phases, MUT-8 thus locally restructures the condensate, which can in turn affect the material and interfacial properties. The double-mutant-cycle analysis indicates that mutations in the scaffold (MUT-16) and client (MUT-8) contribute to phase separation in a largely additive and decoupled fashion. In all four cycles, mutating the scaffold affected MUT-16 partitioning far more than mutating the client. The conservative (R2K) and disruptive (R2A) scaffold substitutions suppressed phase separation to a similar extent. This implies that the loss of the native arginine residues matters more than the identity of the replacement. The coupling free energies remained small throughout, with uncertainties easily bracketing zero in three of the four cycle. The exception was the cycle containing a conservative scaffold mutation (R2K) with a disruptive mutation for the client (Y2A), where we observed only weak coupling.

The active-learning framework efficiently generated peptide clients that progressively strengthened MUT-16 phase separation upon recruitment. After 4 iterations, Δ*G*_trans_(MUT-16) became significantly more negative than the previous iterations. At later iterations, the variants clustered more tightly and showed a growing tendency to self-associate, both predicted and observed. Beyond this point, optimization favors intrinsic self-condensation over scaffold recruitment, marking a practical upper bound. The change in Δ*G*_trans_(MUT-16) was strongly correlated with the conformational properties of MUT-16 at the condensate interface. Specifically, more negative values of Δ*G*_trans_ were associated with increased chain swelling and stronger alignment normal to the interface, consistent with the relationship previously reported by von Bülow *et al.*^35^ At the sequence level, the designed clients shifted toward aromatic residues and positively charged residues, at the expense of negative, hydrophobic, and polar residues. This finding is in agreement with the known sequence gram-mar of phase separation.^10,13^ Active learning thus provides a practical route to designing condensate-modulating peptides while recovering the sequence determinants of phase behavior.

Finally, recruiting wild-type MUT-8 into the condensate markedly altered its material properties. Meanwhile. across iterations, the viscosity increased steadily, whereas the surface tension showed no clear trend despite it being different from the value for wild-type MUT-8. The viscosity correlated strongly with both scaffold partitioning and the interfacial conformational parameters. However, despite systematic variations in the interfacial conformations, the surface tension did not show any sign of correlation.

Since the material properties of biomolecular condensates are increasingly tied to physiological function and to disease-associated aberrant phase transitions,^19^ peptides that bind a chosen scaffold and tune these properties on demand represent a promising approach for developing new therapies. Such designed clients could be used to steer condensate mechanics toward a desired set point without abolishing the condensed state.^55,84^

A natural extension is to move beyond optimizing a single thermodynamic objective and instead train a multi-objective design loop that at the same time optimizes the material properties of interest, such as viscosity or surface tension, rather than only focusing on partitioning as a surrogate.^22^ Coupling this objective with constraints that preserve a balanced residue composition profile would discourage the optimizer from collapsing onto degenerate, compositionally extreme sequences. Such an approach would yield peptides that are both biologically plausible and effective at providing the targeted property.

## Data and Software Availability

Simulation scripts and data related to this publication are available at https://github.com/rodbadr/proteinSimulationsHOOMD4 and https://doi.org/10.5281/zenodo.20827922, respectively.

## Acknowledgement

K.G. was funded by the Deutsche Forschungsgemeinschaft (DFG, German Research Foundation), Project No. 233630050 – TRR 146. This project was funded by SFB 1551 Project No. 464588647 of the DFG (Deutsche Forschungsgemeinschaft). L.S.S. acknowledges support by ReALity (Resilience, Adaptation and Longevity) and Forschungsinitiative des Landes Rheinland-Pfalz. A.C.S and L.S.S. thank M^3^ODEL for support. We gratefully acknowledge the advisory services offered and the computing time granted on the supercomputers Mogon II at Johannes Gutenberg University Mainz, which is a member of the AHRP (Alliance for High-Performance Computing in Rhineland Palatinate) and the Gauss Alliance e.V. We thank Dr. Rene Ketting, N. Ferruz, Dr. Lindorff-Larsen, Dr. Sebastian Falk, L. Baltz and Y. Tuchkov for inspiring discussions.

