## Supplementary material for "Active Learning–Guided Peptide Design for Modulating Condensate Properties upon Recruitment": Supplimentary text and figures

### Supporting Information

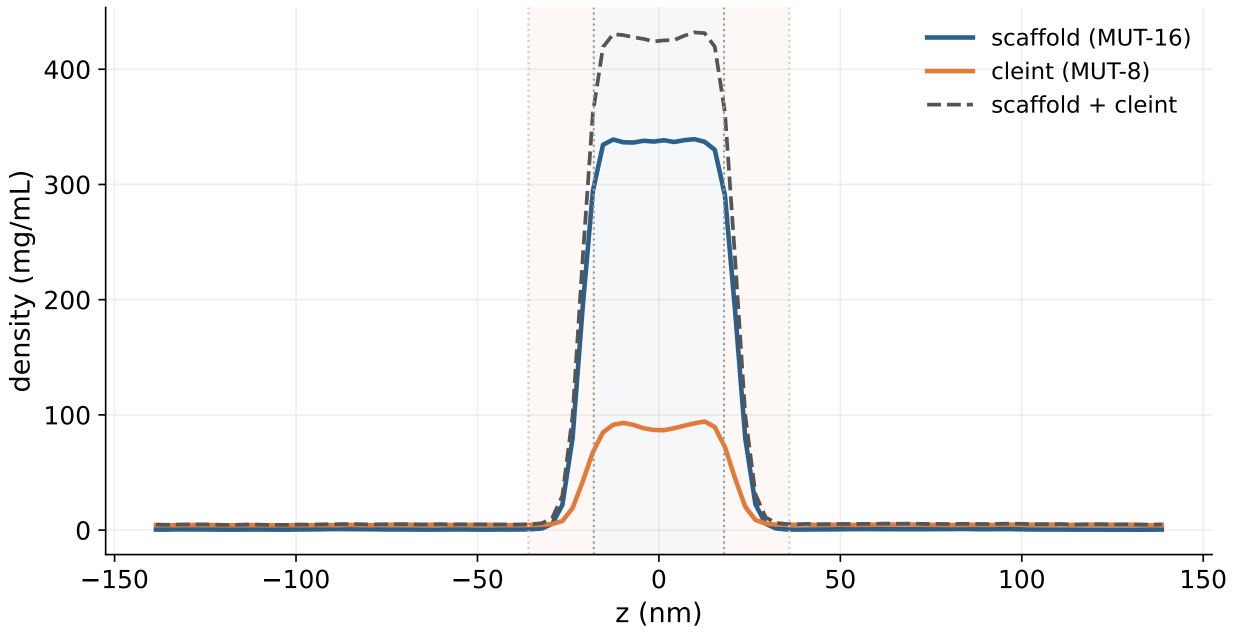


Figure S1: Density profile of scaffold (MUT-16) and client (MUT-8) along the z-dimension of the simulation box.

#### Double-mutant thermodynamic cycle method

To quantify the energetic contribution of Arg–Tyr interactions to MUT-8 recruitment into the MUT-16 condensate, we employed a double-mutant thermodynamic cycle.[72,85,86]

Mutations were introduced at both interaction partners to perturb these contacts: seven Arg residues in MUT-16 M8BR were mutated to Lys and Ala, and eight Tyr residues in the MUT-8 N-terminal domain were mutated to Phe and Ala, enabling both conservative and disruptive substitutions.

The analysis is based on the transfer free energy, $\Delta G_{\mathrm{trans}}$, computed for the wild-type, single-mutant, and double-mutant systems. These values define a thermodynamic cycle from which the coupling free energy is obtained as

$$\Delta\Delta G_{\mathrm{int}} = (\Delta G^{\mathrm{double}}_{\mathrm{trans}} - \Delta G^{\mathrm{Mut1}}_{\mathrm{trans}}) - (\Delta G^{\mathrm{Mut2}}_{\mathrm{trans}} - \Delta G^{\mathrm{WT}}_{\mathrm{trans}}).$$

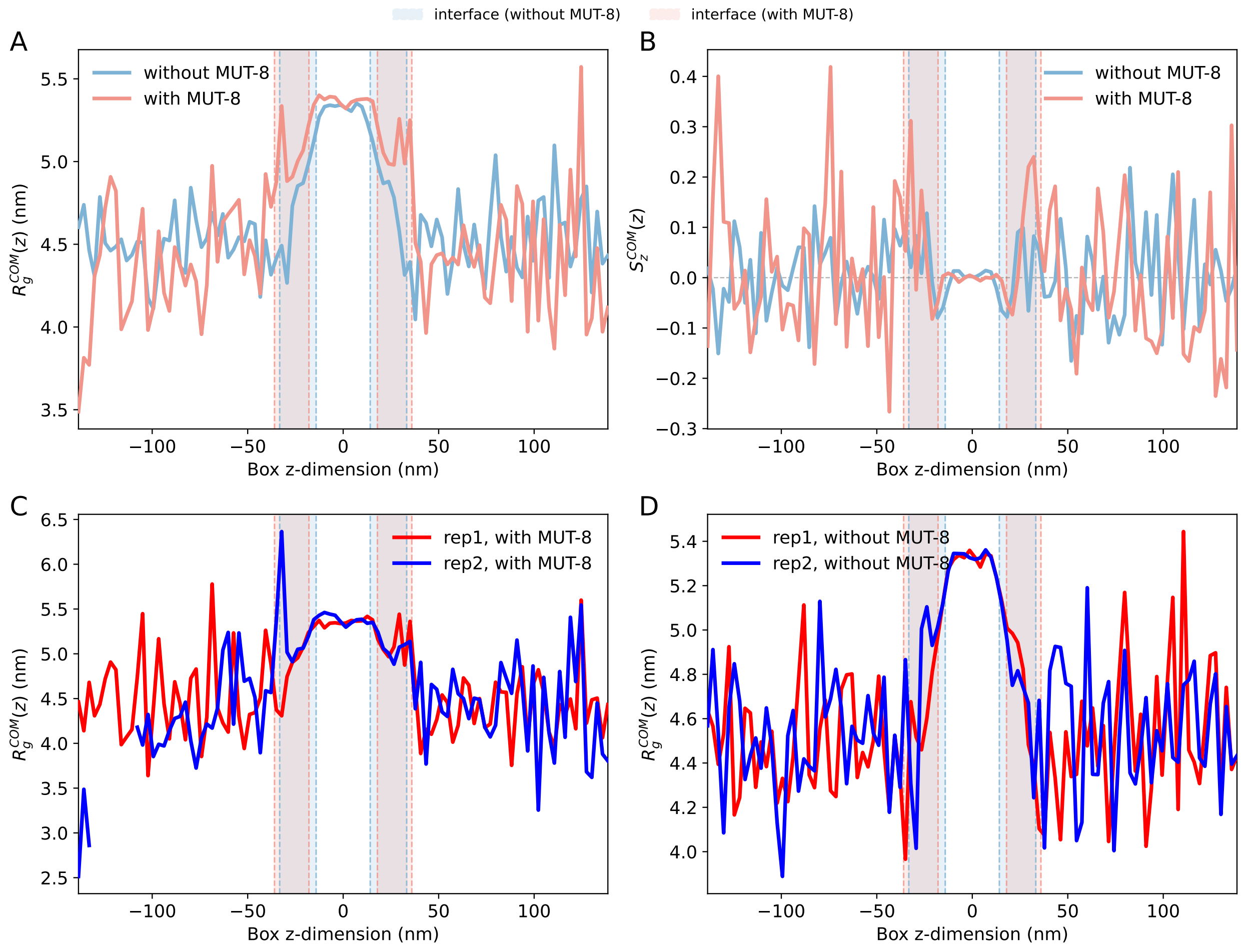


Figure S2: Structural and conformational properties of the MUT-16 condensate with and without MUT-8 peptides. **(A)** Center of mass binned radius of gyration ($R_g$) profile along the $z$-axis, reporting on the size and compactness of individual chains across the slab. **(B)** Center of mass binned orientational order parameter $S_z = \langle P_2(\cos\theta_i)\rangle$ along the $z$-axis, where $\theta_i$ is the angle between the principal axis of chain $i$ (its longest elongation) and the $z$-axis of the simulation box. $S_z$ quantifies chain alignment along $z$: values near 1 indicate strong alignment with the $z$-axis, 0 corresponds to isotropic orientation, and negative values indicate preferential alignment perpendicular to $z$. **(C)** Comparison of the profiles for the two replicas with MUT-8 peptides. **(C)** Comparison of the profiles for the two replicas without MUT-8 peptides.

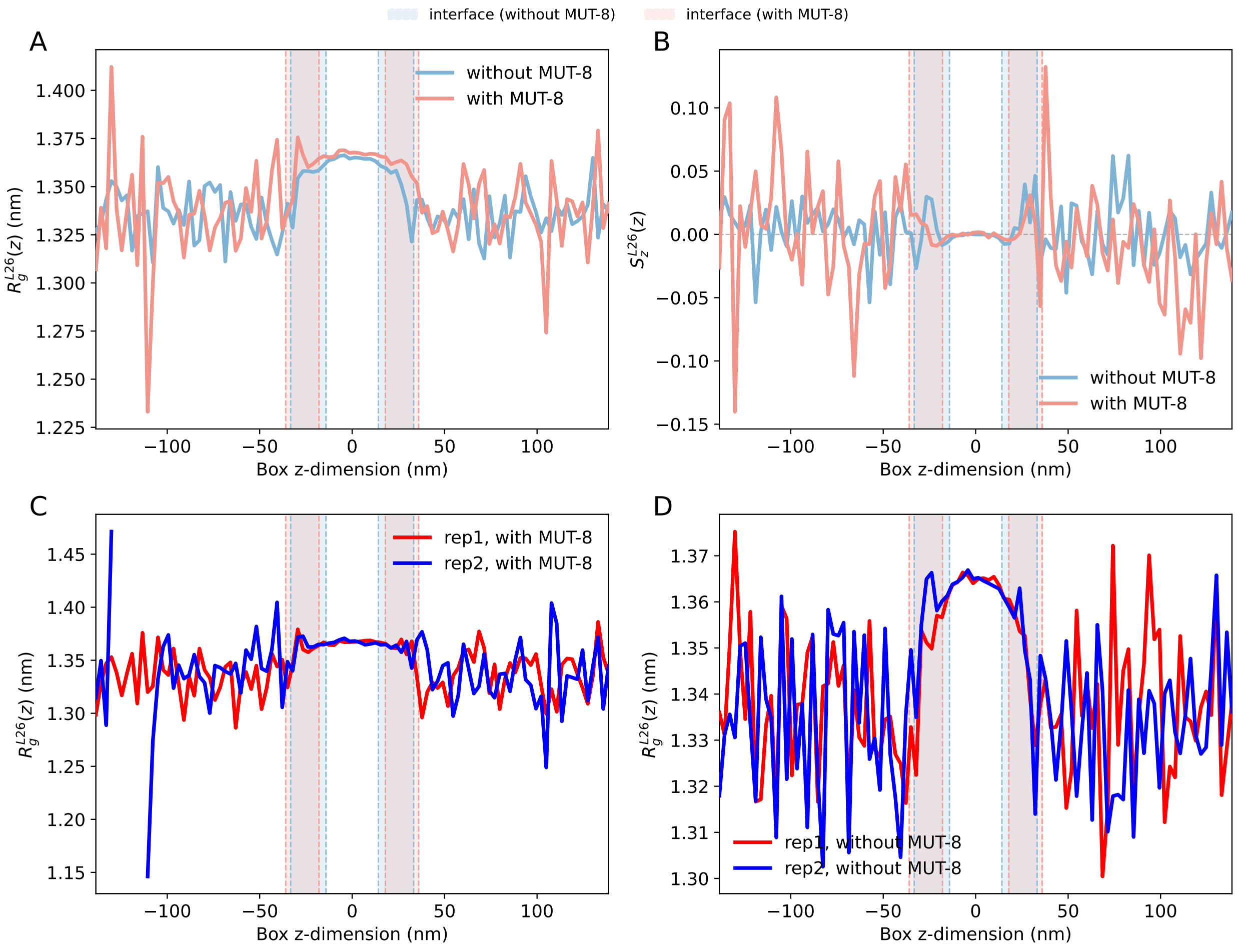


Figure S3: Structural and conformational properties of the MUT-16 condensate with and without MUT-8 peptides, using non-overlapping segments of with length of 26 residues. **(A)** Center of mass binned radius of gyration ($R_g^{l26}$) profile along the $z$-axis, reporting on the size and compactness of individual chains across the slab. **(B)** Center of mass binned orientational order parameter $S_z^{l26} = \langle P_2(\cos\theta_i)\rangle$ along the $z$-axis, where $\theta_i$ is the angle between the principal axis of chain $i$ (its longest elongation) and the $z$-axis of the simulation box. $S_z$ quantifies chain alignment along $z$: values near 1 indicate strong alignment with the $z$-axis, 0 corresponds to isotropic orientation, and negative values indicate preferential alignment perpendicular to $z$. **(C)** Comparison of the profiles for the two replicas with MUT-8 peptides. **(C)** Comparison of the profiles for the two replicas without MUT-8 peptides.

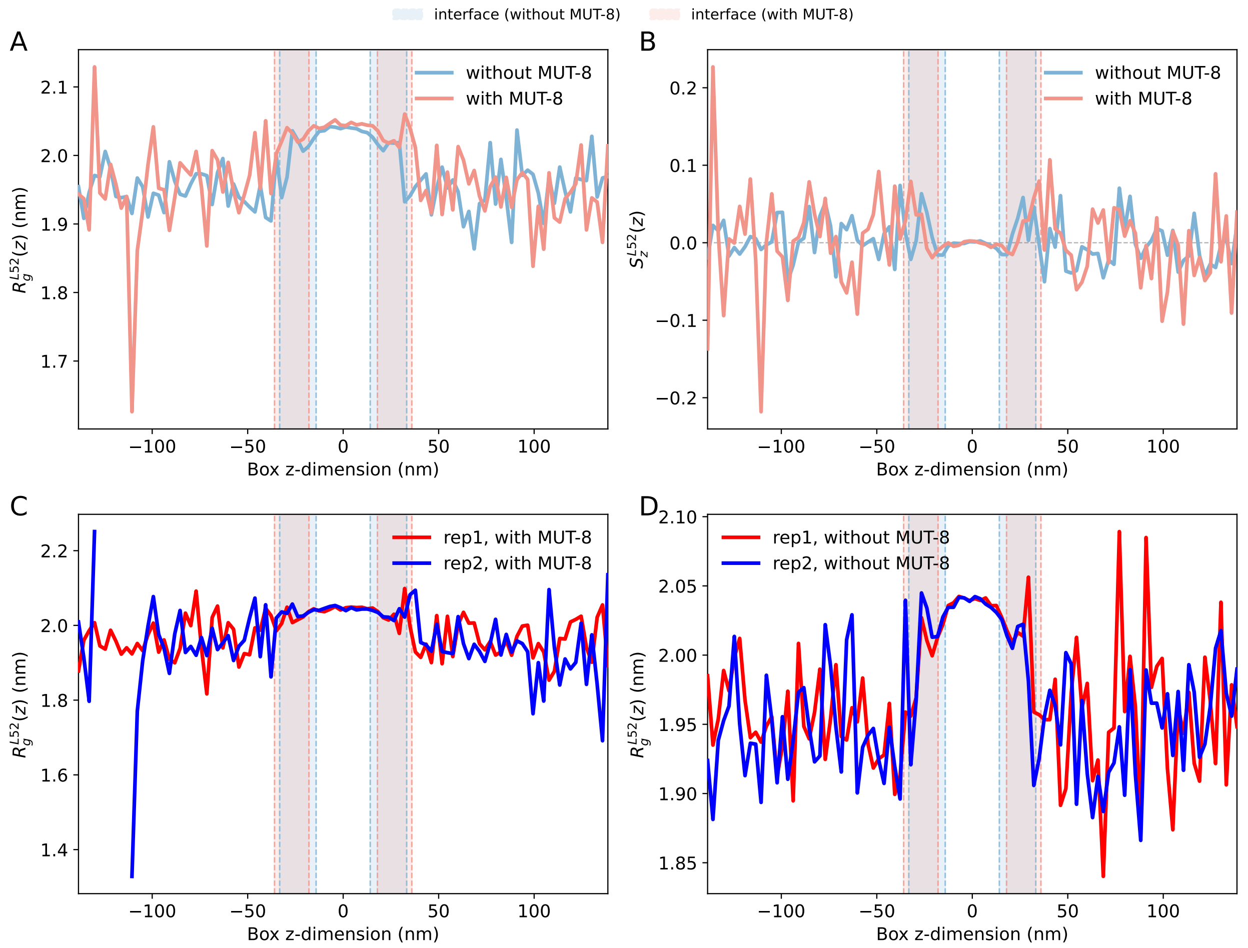


Figure S4: Structural and conformational properties of the MUT-16 condensate with and without MUT-8 peptides, using non-overlapping segments of with length of 52 residues. **(A)** Center of mass binned radius of gyration ($R_g^{l52}$) profile along the $z$-axis, reporting on the size and compactness of individual chains across the slab. **(B)** Center of mass binned orientational order parameter $S_z^{l52} = \langle P_2(\cos\theta_i)\rangle$ along the $z$-axis, where $\theta_i$ is the angle between the principal axis of chain $i$ (its longest elongation) and the $z$-axis of the simulation box. $S_z$ quantifies chain alignment along $z$: values near 1 indicate strong alignment with the $z$-axis, 0 corresponds to isotropic orientation, and negative values indicate preferential alignment perpendicular to $z$. **(C)** Comparison of the profiles for the two replicas with MUT-8 peptides. **(C)** Comparison of the profiles for the two replicas without MUT-8 peptides.

where in order from left to right we have the transfer free energy of the double mutant, the case where only one species was mutated to Mut1, the case where only the other species was mutated to Mut2, and finally the wild type (WT). The coupling free energy $\Delta\Delta G_{\text{int}}$ then measures the difference between the effect of introducing Mut2 to the WT system and introducing it to the Mut1 system. If the two mutations Mut1 and Mut2 are decoupled, then introducing Mut2 should have the same effect regardless of the state of the other species (whether it was mutated or not), resulting in $\Delta G_{\text{trans}}^{\text{double}} \approx 0$. Otherwise, $\Delta G_{\text{trans}}^{\text{double}} \neq 0$ would indicate that the mutations are coupled.

### Double-mutant thermodynamic cycle analysis

We first examined a pair of conservative mutations designed to probe the interaction between the scaffold and client without grossly altering their chemistry. The seven Arg residues in MUT-16 (M8BR+FFR) were substituted with Lys (7R→K, labeled R2K), and the eight Tyr residues in the MUT-8 N-terminal domain were substituted with Phe (8Y→F, labeled Y2F). These substitutions are conservative because the essential chemical features are largely preserved: Lys retains the positive charge of Arg, and Phe retains the aromatic character of Tyr. By combining the two single mutants into a double mutant and comparing the four states through a thermodynamic cycle,[72] we can test whether the Arg–Tyr interaction between scaffold and client contributes additively to MUT-16 phase separation or whether the two perturbations are energetically coupled: a coupling free energy $\Delta\Delta G_{\text{int}}$ near zero indicates additive, independent contributions, whereas a sizeable $\Delta\Delta G_{\text{int}}$ would signal a direct, non-additive interaction. We quantified each state from the MUT-16 density profile along the slab $z$-axis and the corresponding transfer free energy, $\Delta G_{\text{trans}}$(MUT-16).

The density profiles show that the wild-type scaffold/wild-type client combination (MUT-16 WT + MUT-8 WT) exhibits the strongest co-phase separation, with the highest partition coefficients for both scaffold and client (Fig. S5A, Table S1). Here MUT-16 partitions strongly into the dense phase ($K \approx 588.3$; $\rho_{\text{dense}}$(MUT-16) $\approx$ 339.1 mg/ml, $\rho_{\text{dilute}}$(MUT-16) $\approx$

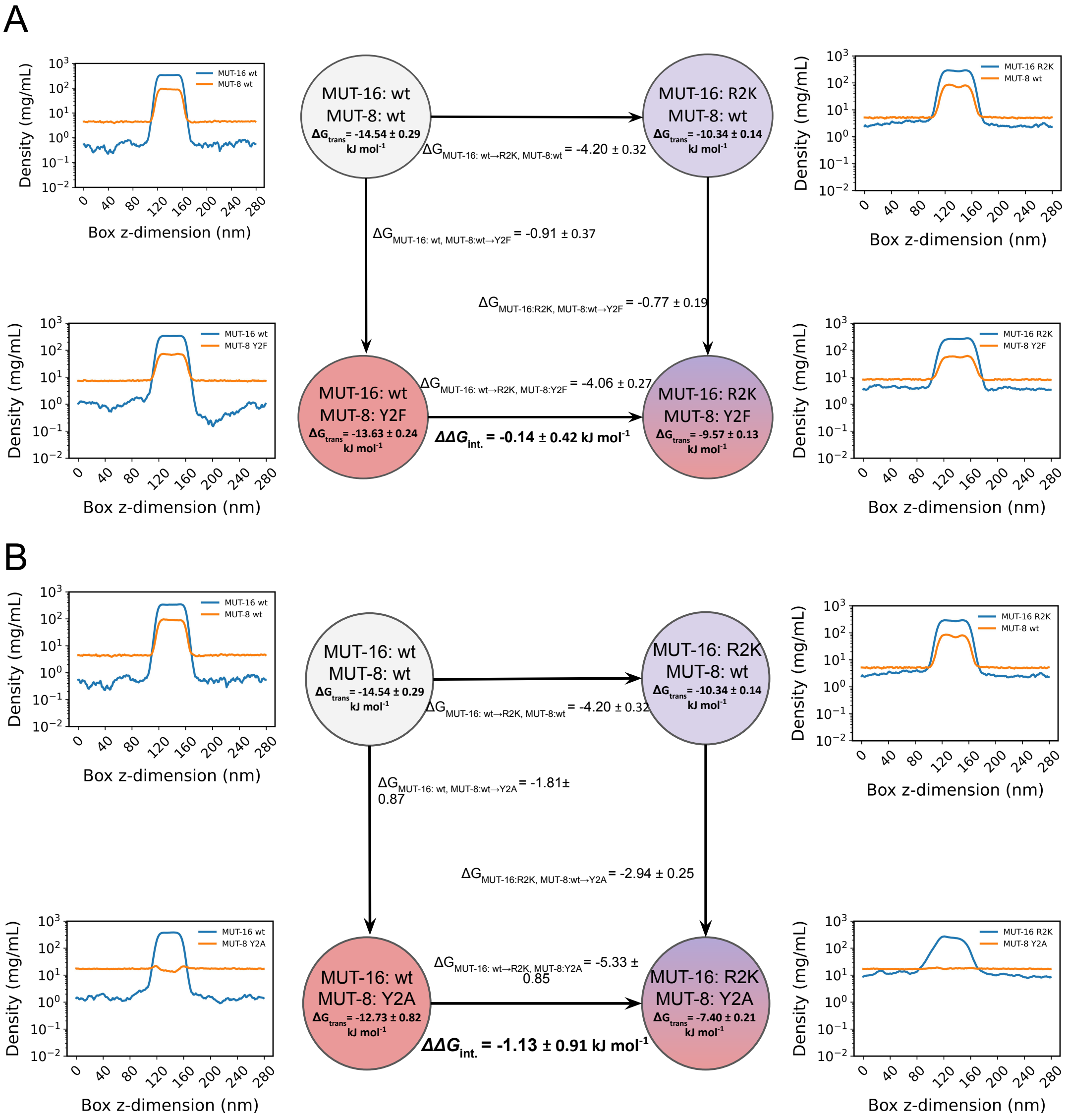


Figure S5: Density profiles and double-mutant thermodynamic cycles quantifying the interaction free energy between MUT-16 and MUT-8 residues. For each simulation, the MUT-16 transfer free energy, $\Delta G_{\text{trans}}(MUT-16)$, is obtained from the MUT-16 density profile along the slab $z$-axis; closing the cycle over the four $\Delta G_{\text{trans}}$(MUT-16) values gives the interaction coupling free energy, $\Delta\Delta G_{\text{int}}$, between the two mutated residues. **A.** MUT-16 R2K / MUT-8 Y2F system. The central thermodynamic cycle connects the four mutational states: wild-type/wild-type, the two single mutants (MUT-16 R2K and MUT-8 Y2F), and the double mutant (MUT-16 R2K + MUT-8 Y2F). Each node is labeled with its corresponding MUT-16 density profile. **B.** MUT-16 R2K / MUT-8 Y2A system, shown as in (A) with states wild-type/wild-type, the two single mutants, and the double mutant (MUT-16 R2K + MUT-8 Y2A).

0.58 mg/ml, $\Delta G_{\text{trans}}$(MUT-16) $\approx -14.53$ kJ/mol). Mutating only the client (MUT-16 wt + MUT-8 Y2F) reduces partitioning modestly ($K \approx 393.9$, $c_{\text{sat}}$(MUT-16) $\approx 0.86$ mg/ml, $\Delta G_{\text{trans}}$(MUT-16) $\approx -13.62$ kJ/mol), whereas mutating only the scaffold (MUT-16 R2K + MUT-8 wt) has a much larger effect, raising the saturation concentration to $c_{\text{sat}}$(MUT-16) $\approx$ 3.07 mg/ml and dropping the partition coefficient to $K \approx 92.9$ ($\Delta G_{\text{trans}}$(MUT-16) $\approx -10.33$ kJ/mol). The double mutant (MUT-16 R2K + MUT-8 Y2F) shows the weakest partitioning ($K \approx$ 66.3, $c_{\text{sat}}$(MUT-16) $\approx 4.01$ mg/ml, $\Delta G_{\text{trans}}$(MUT-16) $\approx -9.56$ kJ/mol). Thus, while perturbing the client weakens scaffold partitioning, the dominant effect comes from mutating the scaffold itself. Closing the thermodynamic cycle gives $\Delta\Delta G_{\text{int}} \approx -0.14 \pm 0.42$ kJ/mol; this near-zero value indicates negligible energetic coupling, so the conservative scaffold and client mutations contribute additively to the progressive loss of MUT-16 phase separation.

We next combined the conservative scaffold mutation (MUT-16 R2K) with a disruptive client mutation (MUT-8 Y2A) (Fig. S5B, Table S1). For the single client mutant (MUT-16 wt + MUT-8 Y2A), $\Delta G_{\text{trans}}$(MUT-16) $\approx -12.73$ kJ/mol, less negative than the corresponding conservative client mutation (MUT-8 Y2F), indicating that the more disruptive Y2A substitution in the client more strongly reduces scaffold partitioning. The double mutant (MUT-16 R2K + MUT-8 Y2A) reduces partitioning further still, with $\Delta G_{\text{trans}}$(MUT-16) $\approx$ $-7.40$ kJ/mol. Closing the thermodynamic cycle yields $\Delta\Delta G_{\text{int}} \approx -1.13 \pm 0.91$ kJ/mol. A vanishing value $\Delta\Delta G_{\text{int}} = 0$ of the coupling energy is outside the the 68% confidence interval, but within the 95% interval. Therefore, we will consider the mutations as weakly coupled.

Furthermore, we examined additional double mutant cycles in which the scaffold carried the disruptive mutation (MUT-16 R2A), combined with either a conservative (MUT-8 Y2F, Fig. S6A) or a disruptive (MUT-8 Y2A, Fig. S6B) client mutation. Interestingly, the disruptive scaffold mutation (MUT-16 R2A; $K \approx 88.5$, $\Delta G_{\text{trans}}$(MUT-16) $\approx -10.14$ kJ/mol) reduced MUT-16 partitioning to nearly the same extent as the conservative scaffold mutation (MUT-16 R2K; $K \approx 92.9$, $\Delta G_{\text{trans}}$(MUT-16) $\approx -10.33$ kJ/mol). This suggests that, for the scaffold, the loss of phase-separation propensity is driven mainly by removing the native Arg residues rather than by the specific identity of the replacement residue, since both the charge-preserving (Lys) and the charge- and bulk-removing (Ala) substitutions yield comparable effects. The coupling free energies were small in both cycles: $\Delta\Delta G_{\text{int}} \approx -0.56 \pm 0.94$ kJ/mol for the conservative client mutation (MUT-16 R2A + MUT-8 Y2F) and $\Delta\Delta G_{\text{int}} \approx -0.82 \pm 1.27$ kJ/mol for the doubly disruptive cycle (MUT-16 R2A + MUT-8 Y2A). In both cases, the uncertainty brackets zero, so these couplings are consistent with additive, essentially independent contributions of the scaffold and client mutations to MUT-16 partitioning.

In conclusion, these double mutant cycles indicate that the scaffold (MUT-16) and client (MUT-8) mutations contribute largely additively to MUT-16 phase separation.Across all four cycles, scaffold mutations exerted a substantially greater influence on MUT-16 partitioning than client mutations. Furthermore, conservative (R2K) and disruptive (R2A) scaffold substitutions produced comparable reductions in phase separation. Within the CALVADOS2 framework, this behavior is consistent with the similar hydropathy ($\lambda$) parameters assigned to Lys and Ala,[41] indicating that the removal of native Arg residues is more important for condensate stability than the chemical identity of the substituting residue. The coupling free energies were small throughout, with uncertainties that bracket zero, pointing to essentially independent contributions.

#### Fine-Tuning of ProtGPT2

All fine-tuning experiments used ProtGPT2, an autoregressive transformer language model built on the GPT-2 architecture and pre-trained on 48 million protein sequences from the UniRef50 database (version 2021_04). The fine-tuning corpus combined condensate-related proteins from the DisProt database (version 2024_12, accessed February 2025) with biomolecular and synthetic condensate entries from the CD-CODE database; after merging and removing exact duplicates, the dataset contained 6,772 sequences, randomly split

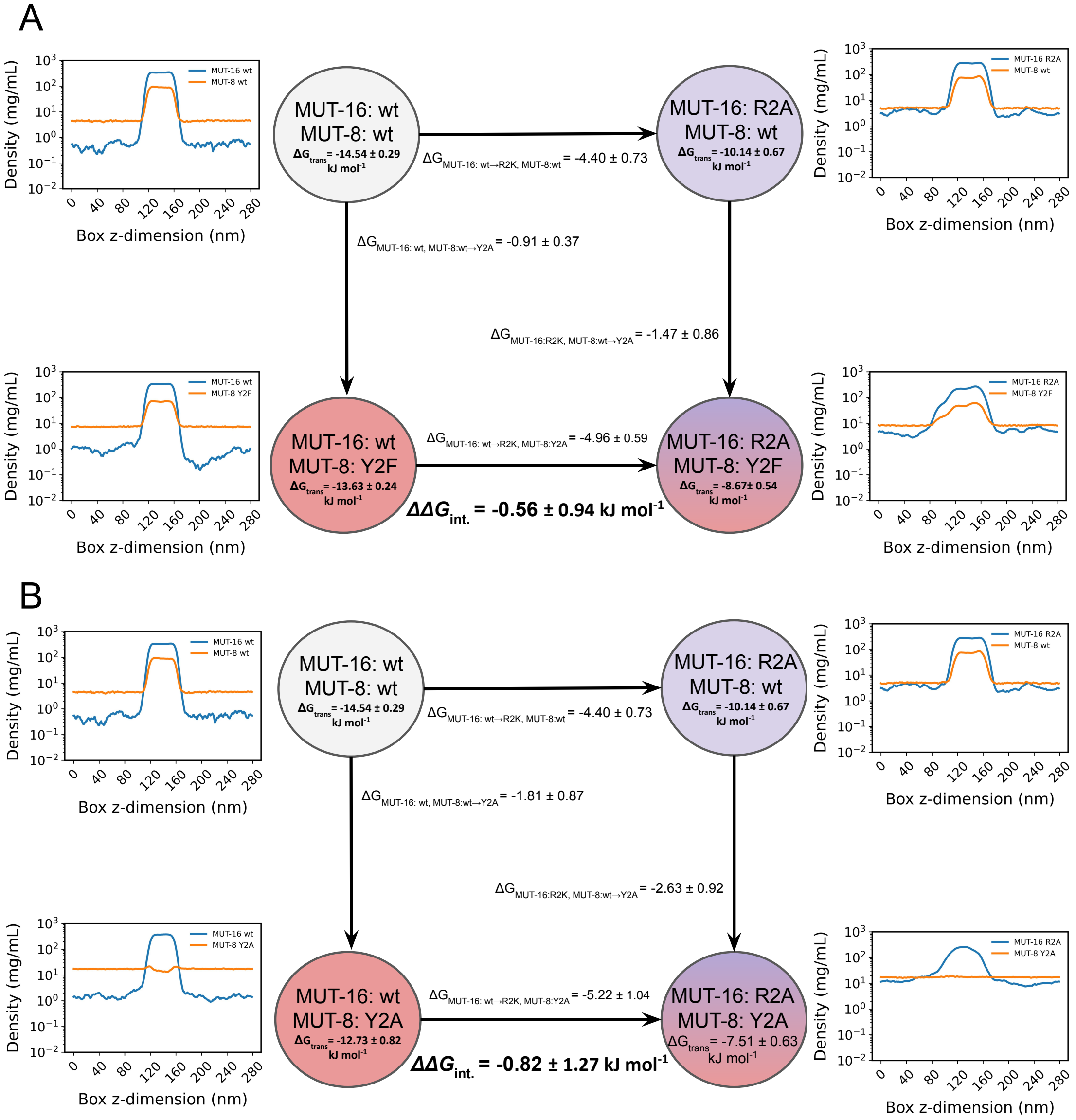


Figure S6: Density profiles and double-mutant thermodynamic cycles used to determine the coupling free energy between the MUT-16 and MUT-8 residues. In each simulation, the MUT-8 transfer free energy, $\Delta G_{\text{trans}}$(MUT-16), is extracted from the MUT-8 density profile along the slab $z$-axis; combining the four $\Delta G_{\text{trans}}$(MUT-16) values around the closed cycle gives the interaction coupling free energy, $\Delta\Delta G_{\text{int}}$, between the two mutated residues. **A.** MUT-16 R2A / MUT-8 Y2F system. The cycle at the center links the four mutational states: wild-type/wild-type, the two single mutants (MUT-16 R2A and MUT-8 Y2F), and the double mutant (MUT-16 R2A + MUT-8 Y2F), with the MUT-16 density profile from each simulation placed beside its node. **B.** MUT-16 R2A / MUT-8 Y2A system, laid out as in (A), spanning wild-type/wild-type, the two single mutants, and the double mutant (MUT-16 R2A + MUT-16 Y2A).

Table S1: Dense- and dilute-phase concentrations, partition coefficient $K$, and transfer free energy $\Delta G_{\text{trans}}(MUT-16)$. Values are mean ± SEM over the replicates.

| Mutation (MUT-16, MUT-8) | $\rho_{\text{dense}}$ (mg/mL) | $\rho_{\text{dilute}}$ (mg/mL) | $K$ | $\Delta G_{\text{trans}}$ (kJ/mol) |
|---|---|---|---|---|
| R2A, Y2A | $269.5 \pm 8.6$ | $10.357 \pm 2.499$ | $27.8 \pm 7.5$ | $-7.505 \pm 0.635$ |
| R2A, Y2F | $237.4 \pm 35.7$ | $5.289 \pm 0.451$ | $45.8 \pm 10.7$ | $-8.665 \pm 0.541$ |
| R2A, WT | $280.2 \pm 0.2$ | $3.444 \pm 0.978$ | $88.5 \pm 25.2$ | $-10.137 \pm 0.668$ |
| R2K, Y2A | $271.6 \pm 18.2$ | $10.713 \pm 1.670$ | $25.7 \pm 2.3$ | $-7.402 \pm 0.205$ |
| R2K, Y2F | $265.6 \pm 1.8$ | $4.018 \pm 0.196$ | $66.3 \pm 3.7$ | $-9.569 \pm 0.127$ |
| R2K, WT | $284.3 \pm 1.7$ | $3.074 \pm 0.208$ | $92.9 \pm 5.8$ | $-10.338 \pm 0.142$ |
| WT, Y2A | $378.4 \pm 2.5$ | $1.523 \pm 0.517$ | $281.3 \pm 97.0$ | $-12.727 \pm 0.821$ |
| WT, Y2F | $337.0 \pm 2.0$ | $0.864 \pm 0.085$ | $393.9 \pm 41.1$ | $-13.628 \pm 0.239$ |
| WT, WT | $339.1 \pm 0.1$ | $0.586 \pm 0.074$ | $588.3 \pm 74.1$ | $-14.538 \pm 0.289$ |

into training (5,417, 80%) and validation (1,355, 20%) sets. Fine-tuning was performed on a single NVIDIA GeForce RTX 4080 GPU using the `run_clm.py` script from Hugging Face, for a maximum of 20 epochs with early stopping on validation loss, a learning rate of $2 \times 10^{-5}$, mixed-precision (FP16) computation, a sequence block size of 256 tokens, and an effective batch size of 8 (per-device batch size of 1 with gradient accumulation over 8 steps). The final model reached a training loss of 3.044 and a validation loss of 6.169 (validation perplexity 477.6); this relatively high perplexity likely reflects the limited dataset size and the narrow nature of intrinsically disordered, condensate-related sequences relative to the broader UniRef50 distribution. The model was therefore used for downstream sequence generation and evaluated on biological-relevance metrics rather than perplexity alone. Comparison of sequences generated before and after fine-tuning confirms that the procedure shifts the model toward the target sequence space (Figure S7): the median predicted disorder score[87] increases from 0.467 to 0.533, and the amino acid composition shifts toward disorder-promoting residues (notably A, G, P, and S) at the expense of order-promoting and aromatic residues (F, W, Y, I, and C).

In addition to generating 51 amino acid sequences, we wanted to assess the ability of the fine-tuned model to generate longer, continuous IDR segments, which could be important for bio-synthetic applications. Hence, we generated sequences of 95–115 amino acids and characterized two complementary metrics, the length of the longest disordered segment per sequence, and the lengths of all disordered segments pooled across sequences. Before fine-tuning, sequences contained multiple short, fragmented disordered regions interspersed with folded segments. The longest disordered segment per sequence had a median of 53 amino acids whereas the after fine-tuning approach resulted in a median of 111 amino acids, indicating that nearly the entire sequence consists of a single continuous disordered block (Fig. S7D). This was further supported by the distribution of all disordered segments. Before fine-tuning, disordered segments were heterogeneous in length with a median of 49 amino acids, consistent with fragmented disorder. After fine-tuning, the median segment length was 110 amino acids, confirming that disorder is no longer fragmented but spans the full sequence length (Fig. S7 E). Thus, fine-tuning didn't just produce more disordered residues, but rather it reorganized the generated sequences with fewer disordered segment and longer continuous disordered domains.

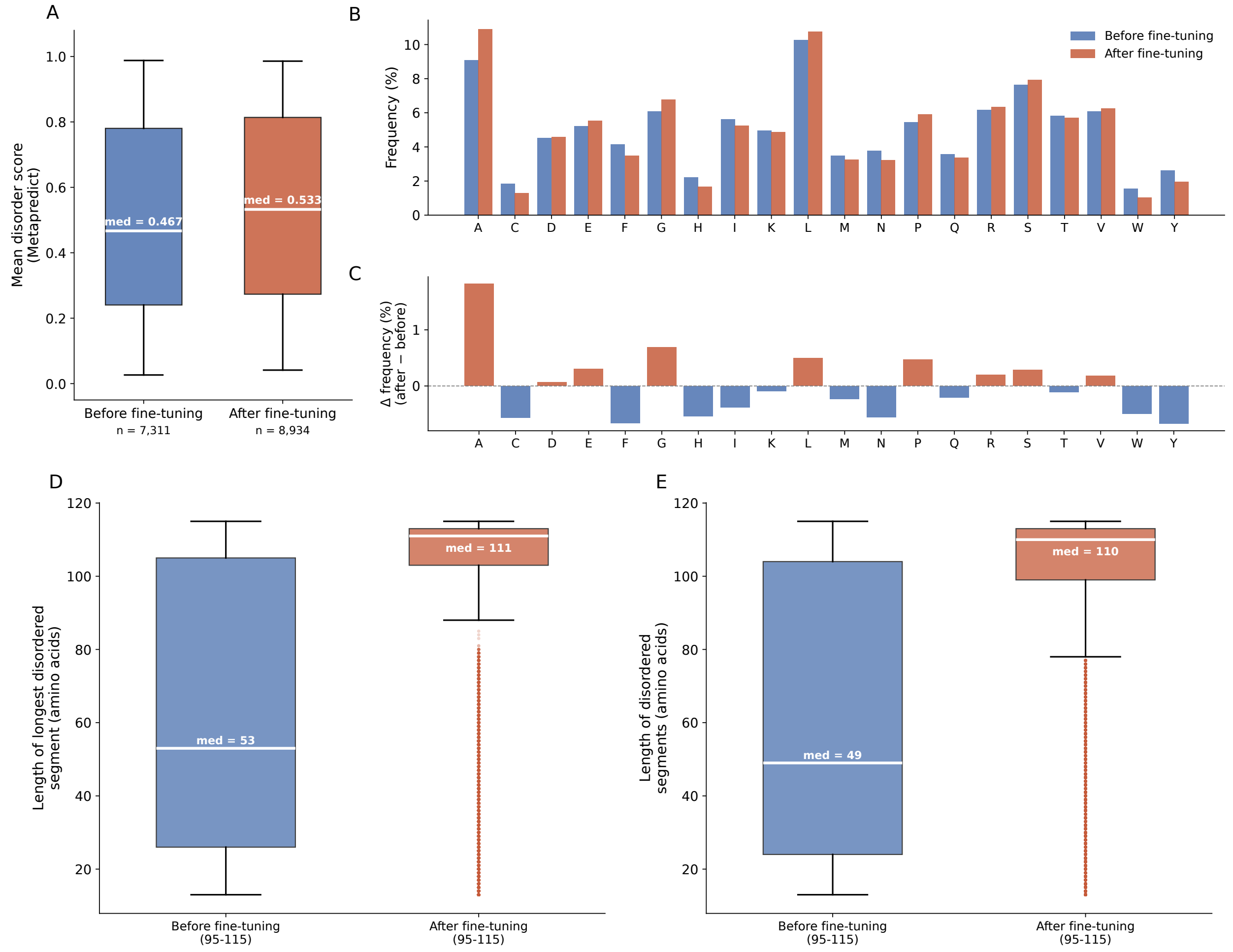


Figure S7: Effect of fine-tuning ProtGPT2[59] on generated sequence properties. **(A)** Distribution of mean per-residue disorder scores (Metapredict[87]) for sequences generated before and after fine-tuning. Fine-tuning shifts the median disorder score upward, consistent with enrichment in intrinsically disordered, condensate-related sequences (sequences of 51 amino acid length). **(B)** Amino acid composition (frequency, %) of generated sequences (sequences of 51 amino acid length) before and after fine-tuning. **(C)** Compositional shift after fine-tuning, shown as the change in frequency for each amino acid ($\Delta = \text{after} - \text{before}$) (sequences of 51 amino acid length). **(D)** Length of the largest disordered segments for peptides of length 95-115 aa as computed by Metapredict[87]) after fine tuning, compared to ProtGPT2 results before finetuning. **(E)** Length of disordered segments peptides of length 95-115 after fine tuning, compared to ProtGPT2 results before finetuning.

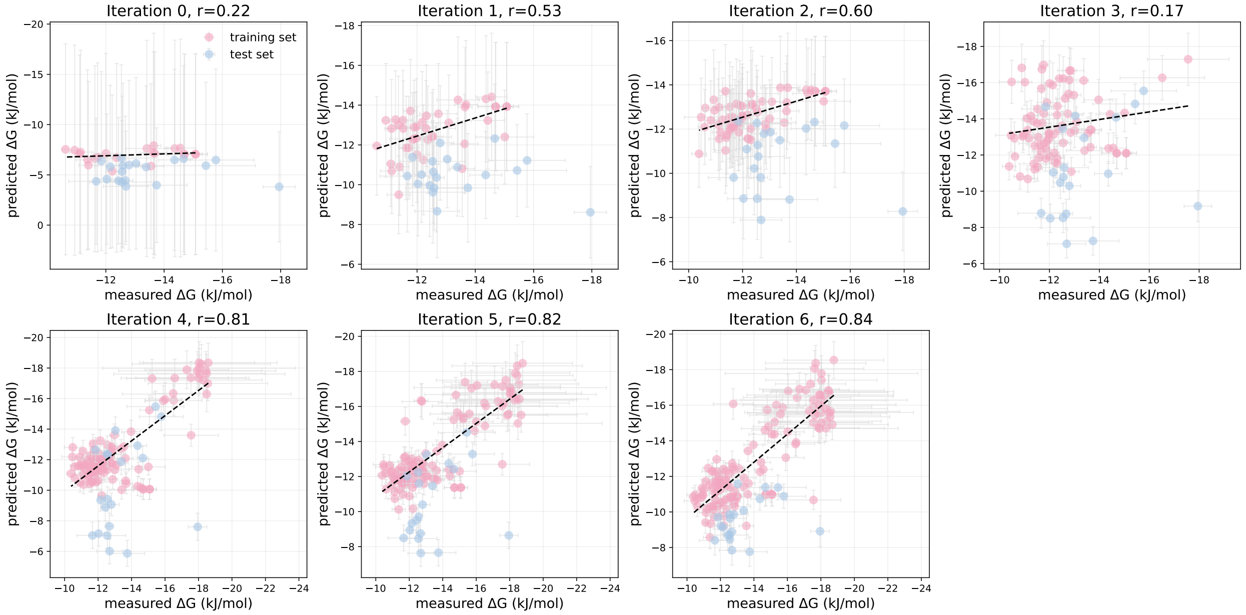


Figure S8: Correlation between predicted and measured values of the transfer free energy $\Delta G_{\mathrm{trans}}$ for peptide variants recruited into the MUT-16 condensate, shown for each iteration of the active learning loop. Predicted values are obtained from the surrogate model (Fig.1) and measured values from MD simulations. Pink points denote the training set, comprising peptide variants accumulated over successive iterations; the correlation coefficient between predicted and measured values increases progressively across iterations, reflecting the improving accuracy of the trained model as more data are added. Blue points denote an independent test set of 20 variant sequences generated by the fine-tuned ProtGPT2 model, overlaid on the same axes to assess how well the model generalises to sequences drawn from a different generative source. The proximity of both sets to the fitted regression line indicates that the model reliably predicts $\Delta G_{\mathrm{trans}}$ for the ProtGPT2-generated variants as well as for the iteratively sampled training sequences.

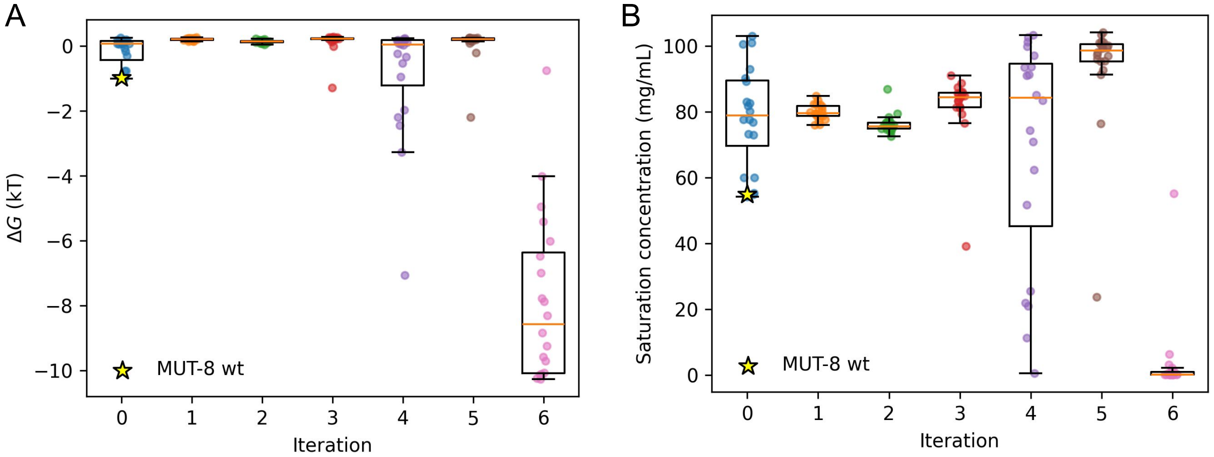


Figure S9: Prediction of phase separation propensity for MUT-8 sequence variants. **A.** Binding free energy ($\Delta G$) of peptide variants generated across successive `wazy`[47] iterations, as calculated by the neural network predictor of von Bülow *et al.*[40] **B.** Predicted saturation concentration of the corresponding MUT-8 mutants obtained over the same `wazy`[47] iterations, reflecting their relative phase separation propensity.

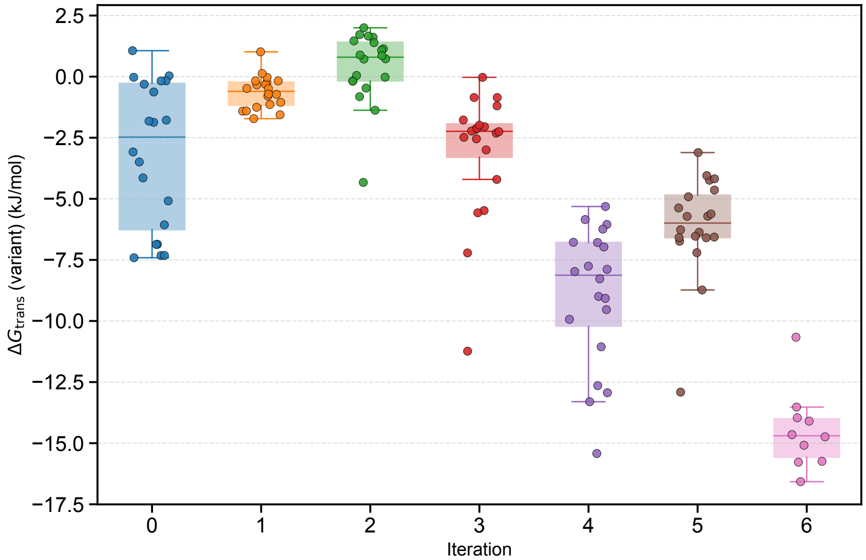


Figure S10: Distribution of $\Delta G_{trans}$(MUT-8) values across successive iterations, computed from the molecular dynamics simulations of client-scaffold interactions. Boxes span the interquartile range (IQR) with the median marked by the central line; whiskers extend to 1.5× IQR. Individual measurements are overlaid as scatter points.

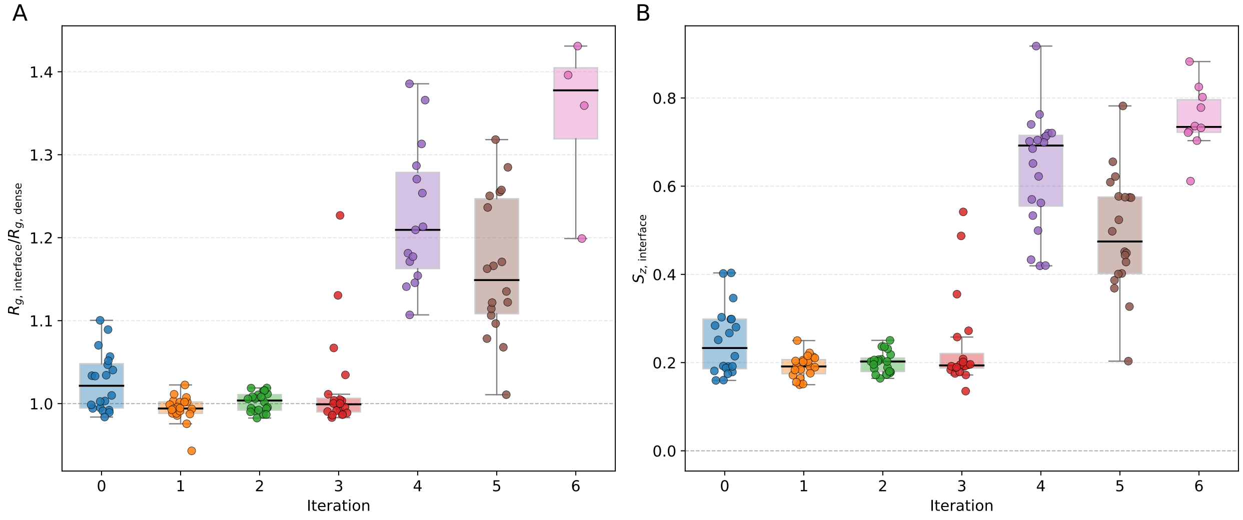


Figure S11: Evolution of MUT-16 interfacial conformational properties across successive peptide variant design iterations obtained from the peaks of the bead-weighted profiles. **(A)** Ratio of the radius of gyration of MUT-16 chains at the interface to that in the dense phase, ($R_{g,\text{interface}}/R_{g,\text{dense}}$). **(B)** Orientational order parameter of MUT-16 chains at the interface, $S_{z,\text{interface}}$.

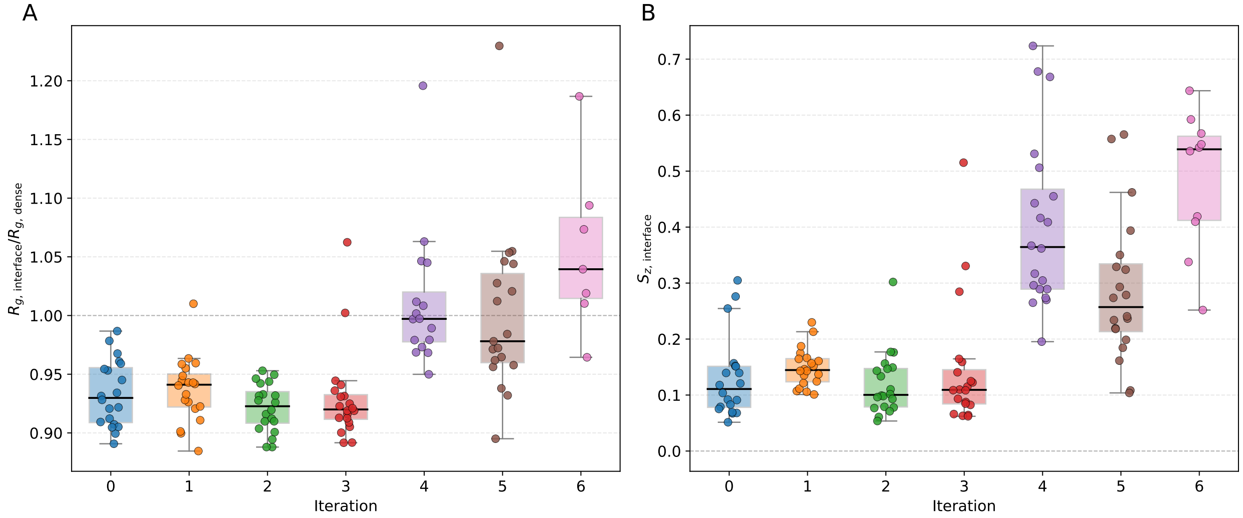


Figure S12: Evolution of MUT-16 interfacial conformational properties across successive peptide variant design iterations obtained from the peaks of the center of mass binned profiles. **(A)** Ratio of the radius of gyration of MUT-16 chains at the interface to that in the dense phase, ($R_{g,\text{interface}}/R_{g,\text{dense}}$). **(B)** Orientational order parameter of MUT-16 chains at the interface, $S_{z,\text{interface}}$.

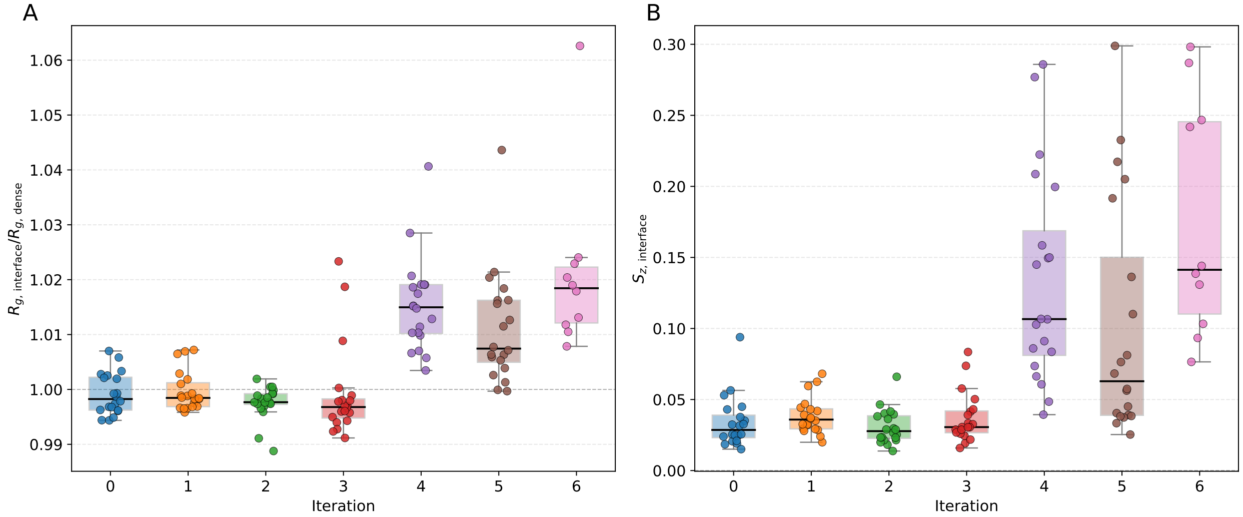


Figure S13: Evolution of MUT-16 interfacial conformational properties across successive peptide variant design iterations for segments composed of 26 residues. **(A)** Ratio of the radius of gyration of MUT-16 chains at the interface to that in the dense phase, ($R_{g,\text{interface}}/R_{g,\text{dense}}$). **(B)** Orientational order parameter of MUT-16 chains at the interface, $S_{z,\text{interface}}$.

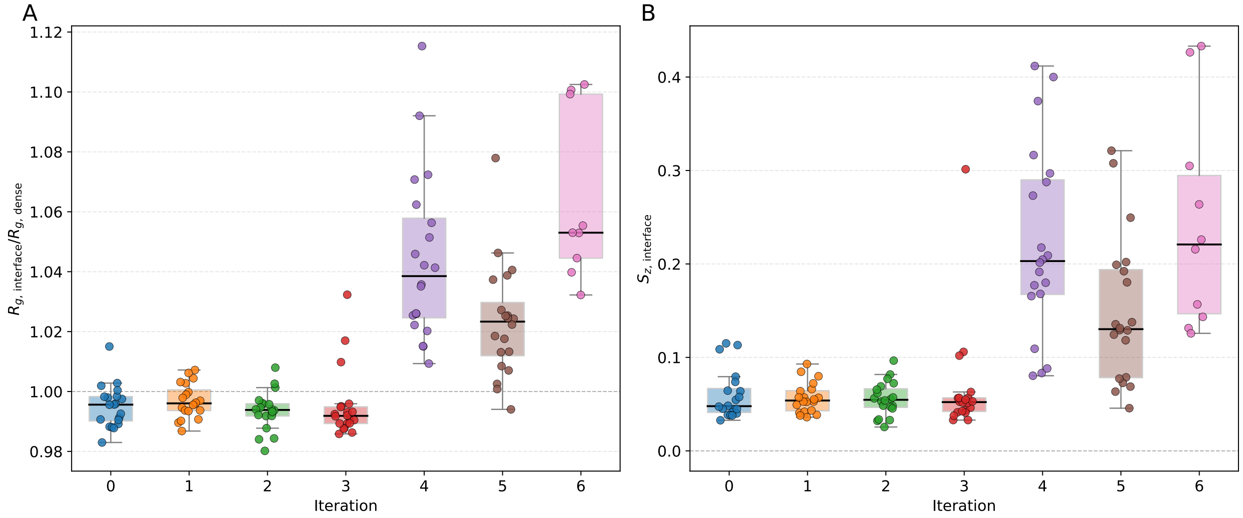


Figure S14: Evolution of MUT-16 interfacial conformational properties across successive peptide variant design iterations for segments composed of 52 residues. **(A)** Ratio of the radius of gyration of MUT-16 chains at the interface to that in the dense phase, ($R_{g,\text{interface}}/R_{g,\text{dense}}$). **(B)** Orientational order parameter of MUT-16 chains at the interface, $S_{z,\text{interface}}$.

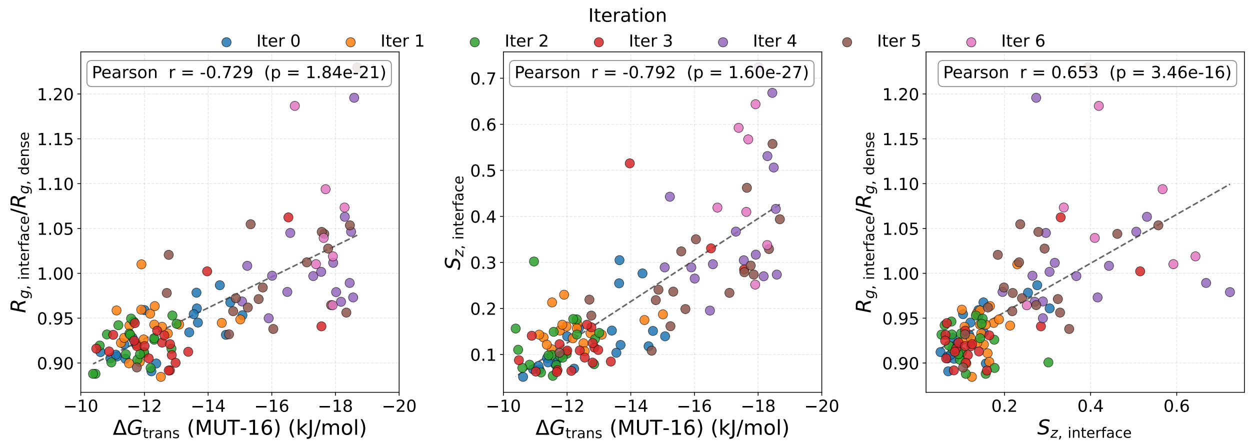


Figure S15: Correlations between the interfacial conformational properties of MUT-16 chains from center of mass binning and the transfer free energy $\Delta G_{\text{trans}}$(MUT-16), evaluated across peptide variants from successive design iterations. The ratio $R_{g,\text{interface}}/R_{g,\text{dense}}$ reports how much MUT-16 chains expand or contract at the interface relative to the condensed interior, $S_{z,\text{interface}}$ is the orientational order parameter of interfacial chains, and $\Delta G_{\text{trans}}$(MUT-16) quantifies the preference of MUT-16 for the dense over the dilute phase. **(A)** $R_{g,\text{interface}}/R_{g,\text{dense}}$ vs. $\Delta G_{\text{trans}}$(MUT-16). **(B)** $S_{z,\text{interface}}$ vs. $\Delta G_{\text{trans}}$(MUT-16), relating interfacial chain alignment to the transfer free energy. **(C)** $R_{g,\text{interface}}/R_{g,\text{dense}}$ vs. $S_{z,\text{interface}}$, relating chain expansion to orientational ordering at the interface. Each point is a distinct peptide variant from the iterative design procedure; the Pearson correlation coefficient is shown in each panel.

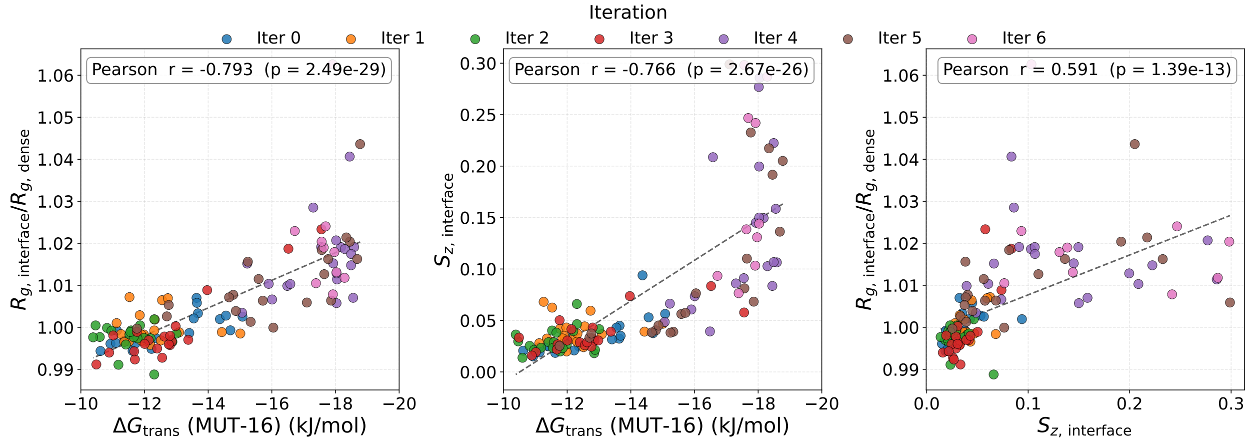


Figure S16: Same as Fig. S15 but for MUT-16 segments composed of 26 residues

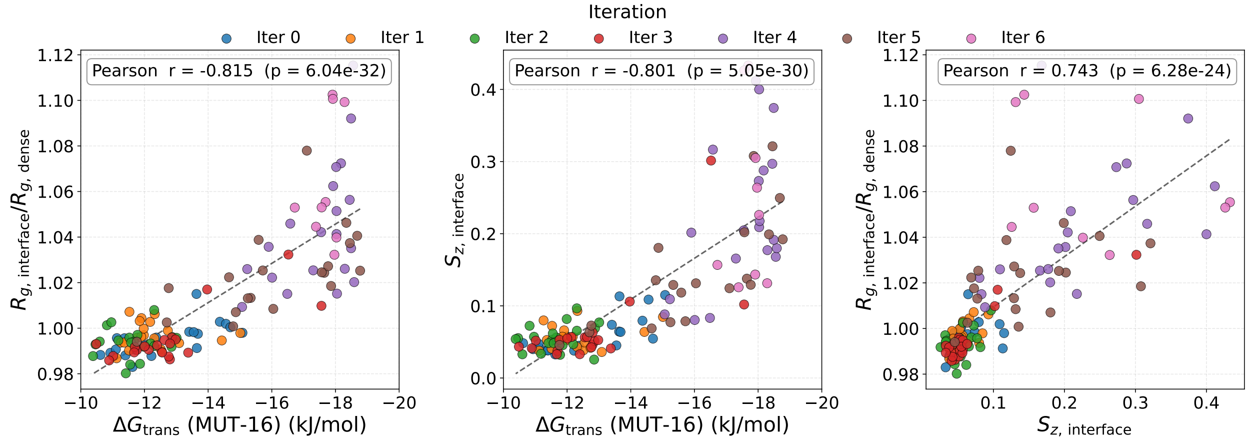


Figure S17: Same as Fig. S15 but for MUT-16 segments composed of 52 residues

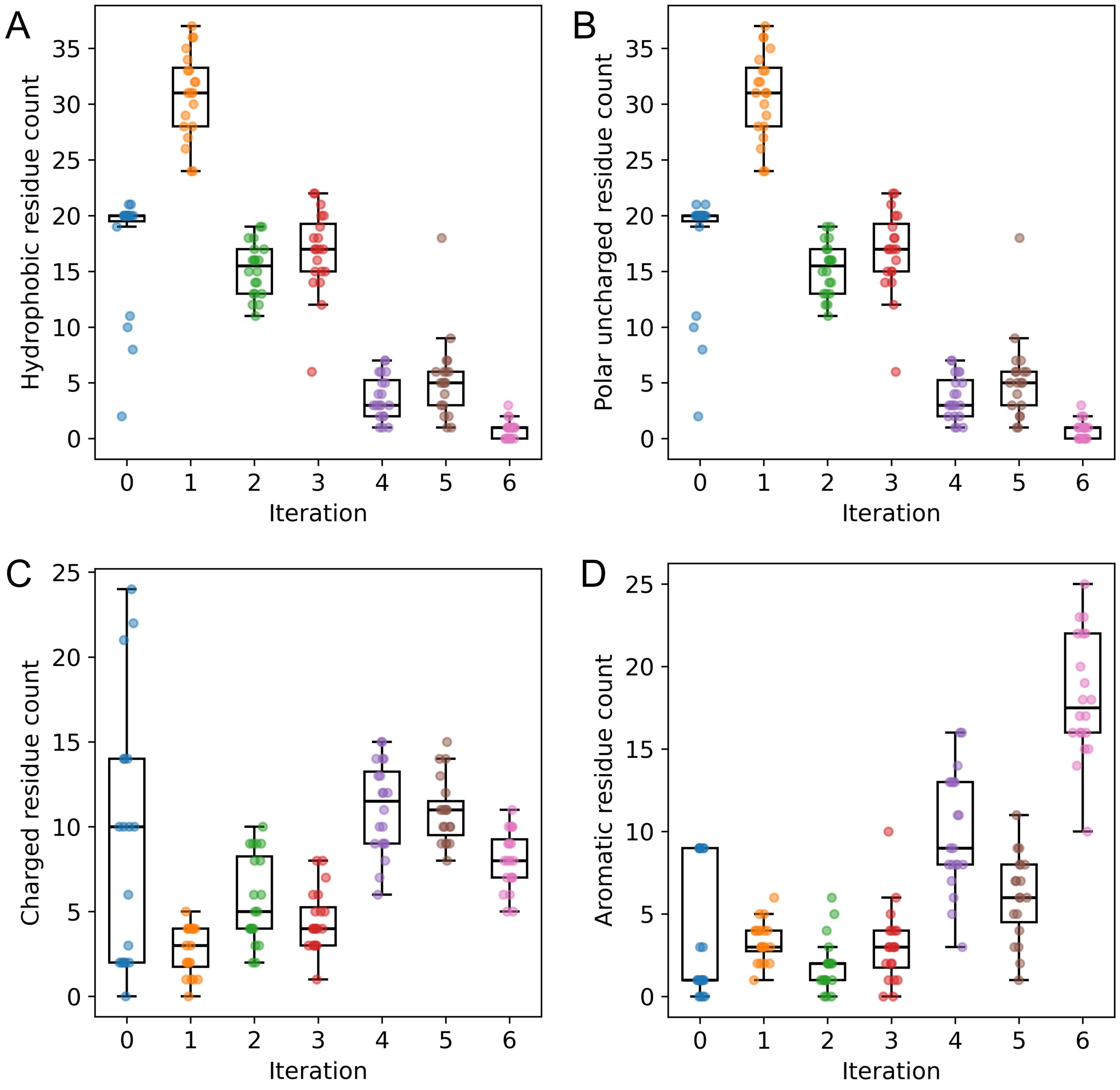


Figure S18: Box-and-whisker plots showing the distribution of residue counts per sequence across iterations. In all panels, the central line represents the median, the box denotes the interquartile range (IQR), whiskers extend to 1.5× IQR, and individual sequences are overlaid as scatter points. **A.** Hydrophobic residues: Ala (A), Val (V), Ile (I), Leu (L), Met (M), Phe (F), Trp (W), and Tyr (Y). **B.** Polar uncharged residues: Ser (S), Thr (T), Asn (N), and Gln (Q). **C.** Charged residues: Lys (K) and Arg (R) (positively charged), and Asp (D) and Glu (E) (negatively charged). **D.** Aromatic residues: Phe (F), Trp (W), and Tyr (Y).

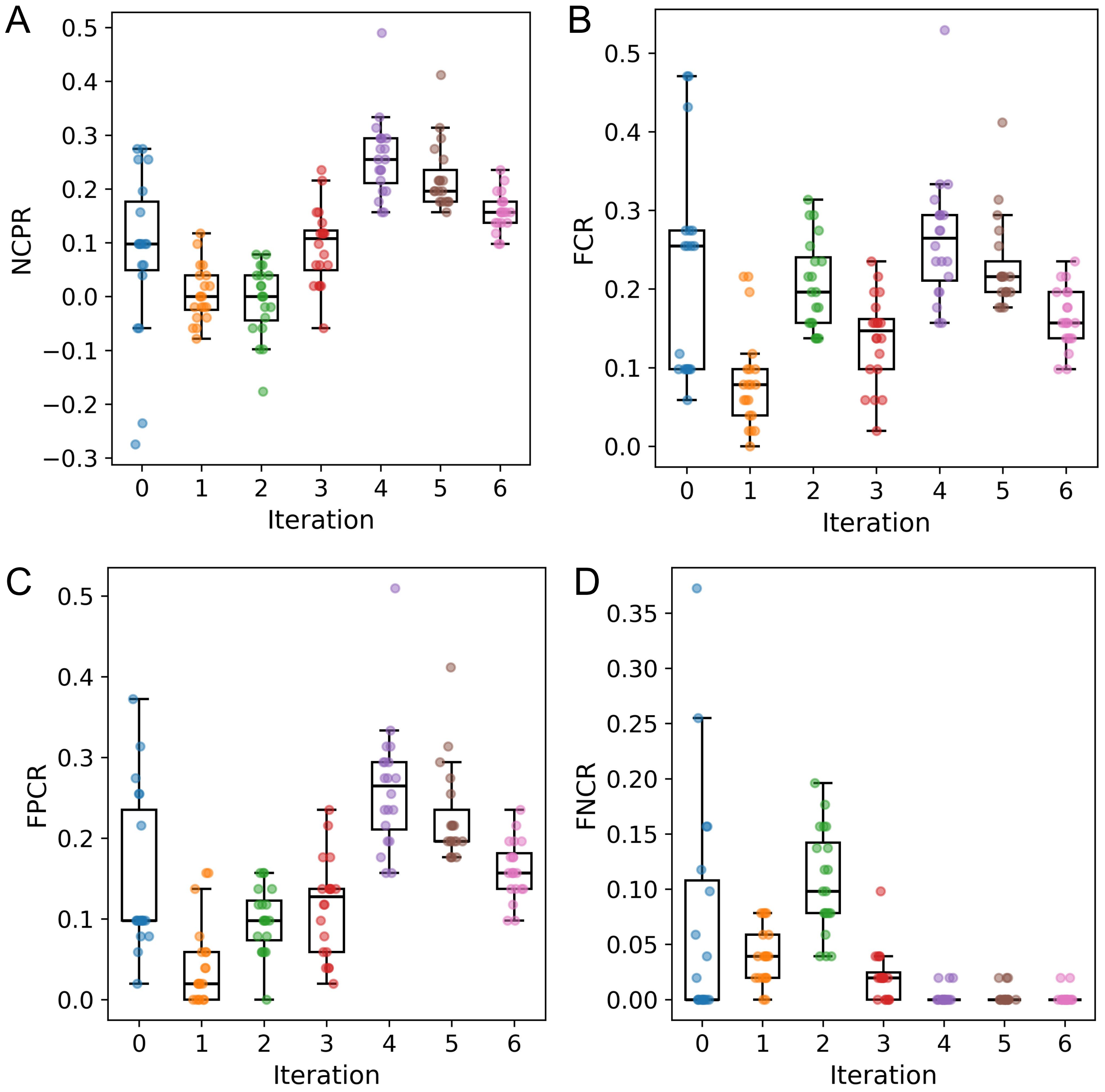


Figure S19: Evolution of charge-related sequence features over active learning iterations. **A.** Net charge per residue (NCPR). **B.** Fraction of charged residues (FCR). **C.** Fraction of positively charged residues (FPCR). **D.** Fraction of negatively charged residues (FNCR). In each panel, box-and-whisker plots show the median and interquartile range, with individual sequence values overlaid as scatter points to illustrate variability within each iteration.

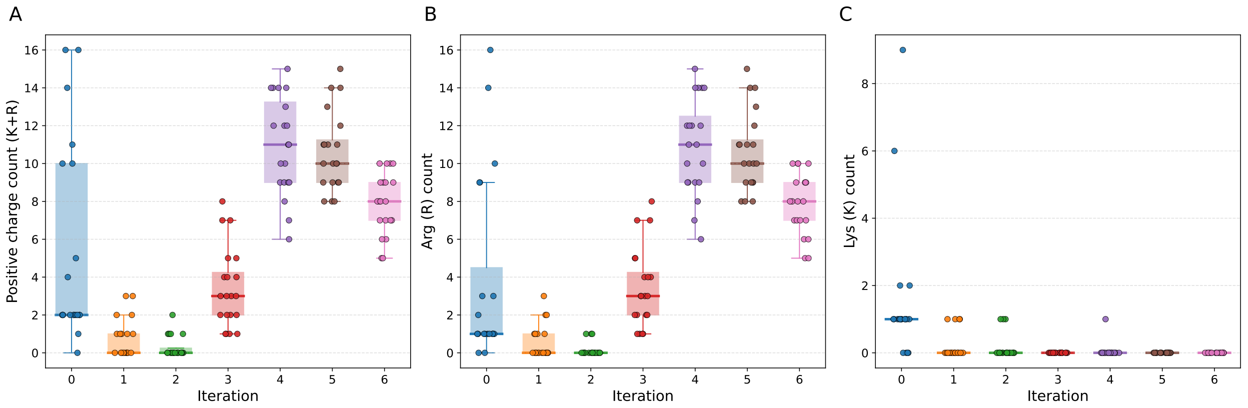


Figure S20: Positively charged residue counts per sequence across design iterations (box plots with overlaid per-mutant points). **(A)** Combined lysine and arginine count (K + R). **(B)** Arginine (R) count. **(C)** Lysine (K) count. Colors denote the iteration.

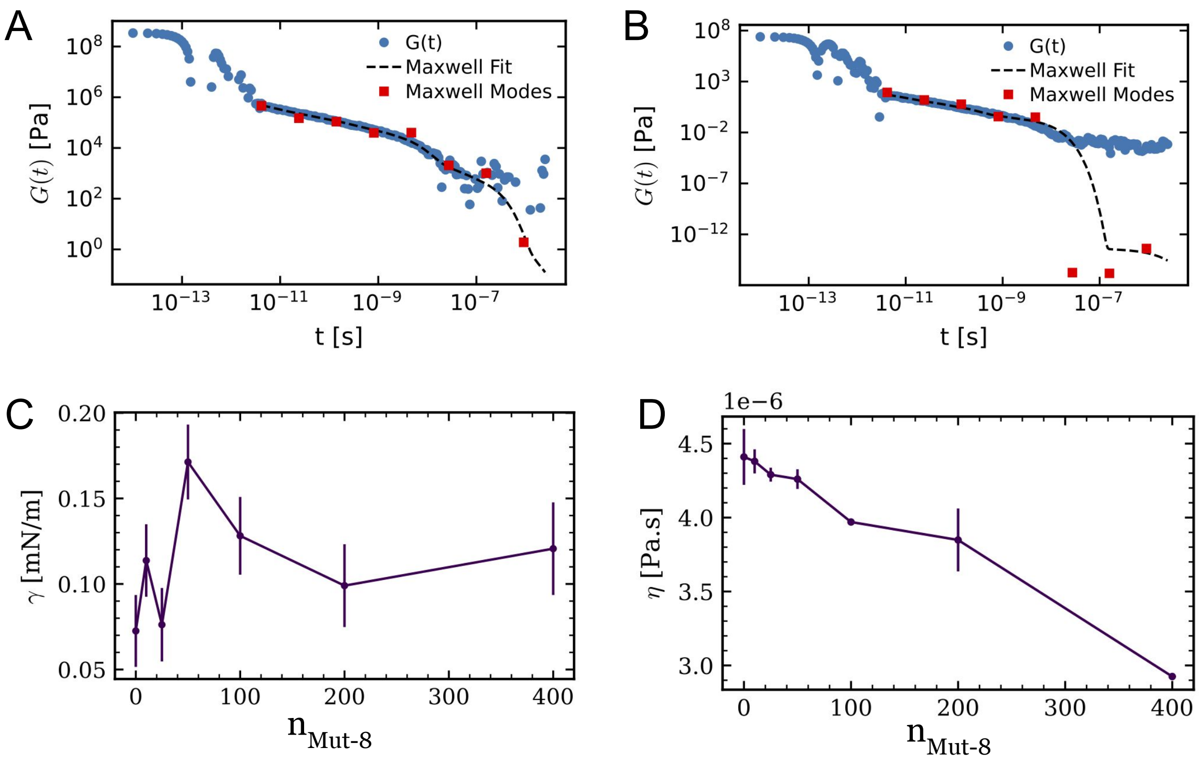


Figure S21: **A.** Stress relaxation of MUT-16 condensates in the absence of MUT-8 peptides. **B.** Stress relaxation of MUT-16 condensates in the presence of 50 MUT-8 peptides. In both cases, Maxwell mode fits are shown for $t \geq 4 \times 10^{-12}$ s. **C.** Surface tension of the condensate as a function of the number of MUT-8 N-terminal PLD peptide chains in the simulation box. **D.** Viscosity of the condensate as a function of the number of MUT-8 N-terminal PLD peptide chains in the simulation box.

#### Statistics of the surface tension calculation

As mentioned in the main text, the surface tension was calculated using the stress tensor anisotropy. We obtain the stress tensor components from HOOMD-blue at runtime. We print values every 0.3ns in simulation time for a total of $10^5$ measurements. Here, we present some relevant statistics. Figure S22 shows the effect of the sample size on the average, error, and distribution of surface tension values. Is it clear that $10^3$ measurements does not result in good statistics. While this improves for a sample size of $10^4$, it is only after $10^5$ that we converge to a proper normal distribution. Figure S23 shows the results of block analysis and the autocorrelation function for $10^5$ surface tension measurements. Both figures show that the values are not correlated, as made clear by the independence of the error on the block size and the immediate decay of the autocorrelation function.

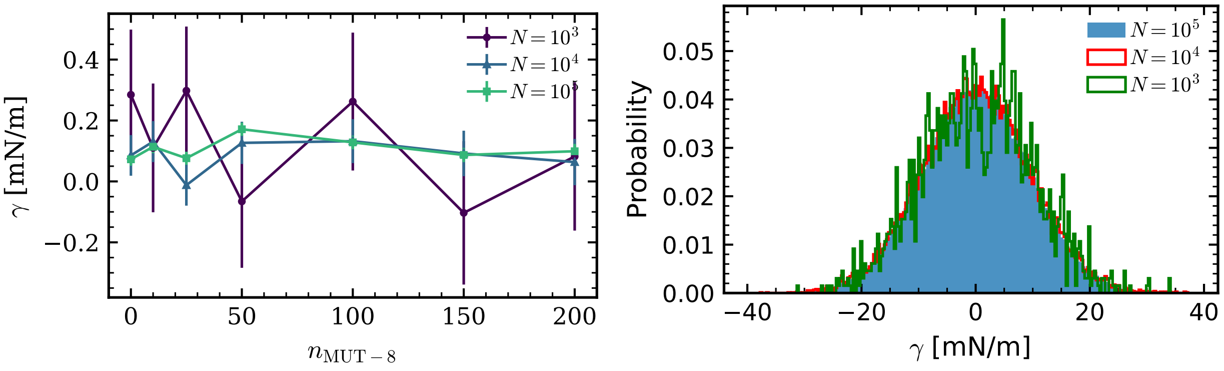


Figure S22: (a) Average surface tension versus the number of WT MUT-8 peptides for different sample sizes as indicated in the legend. (b) Probability distribution for the values of $\gamma$ for different sample sizes for a condensate with no peptides.

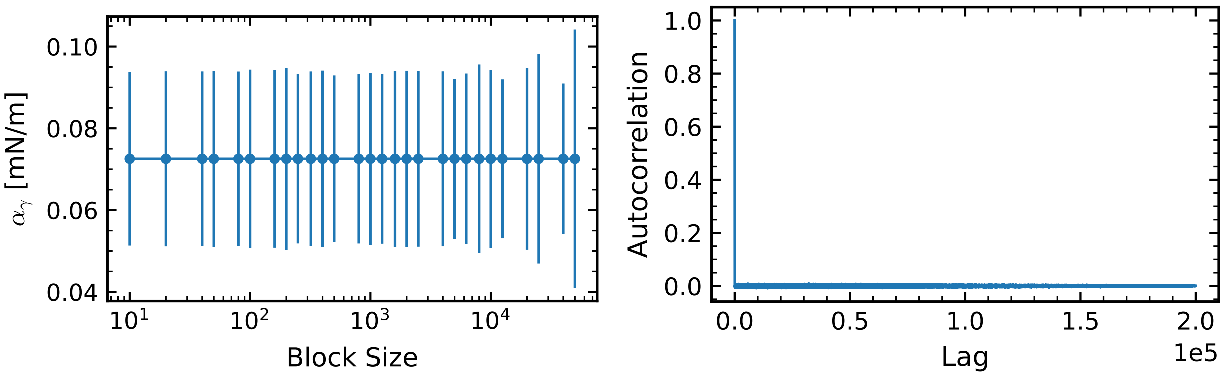


Figure S23: (for SI) WT MUT-16 condensate at $T = 275\ K$. (a) Mean surface tension with error based on block averaging for one of the replicas. (b) Autocorrelation function of the surface tension values. Both figures indicate very small correlations between measurements and that straightforward averaging and calculation of errors is sufficient.

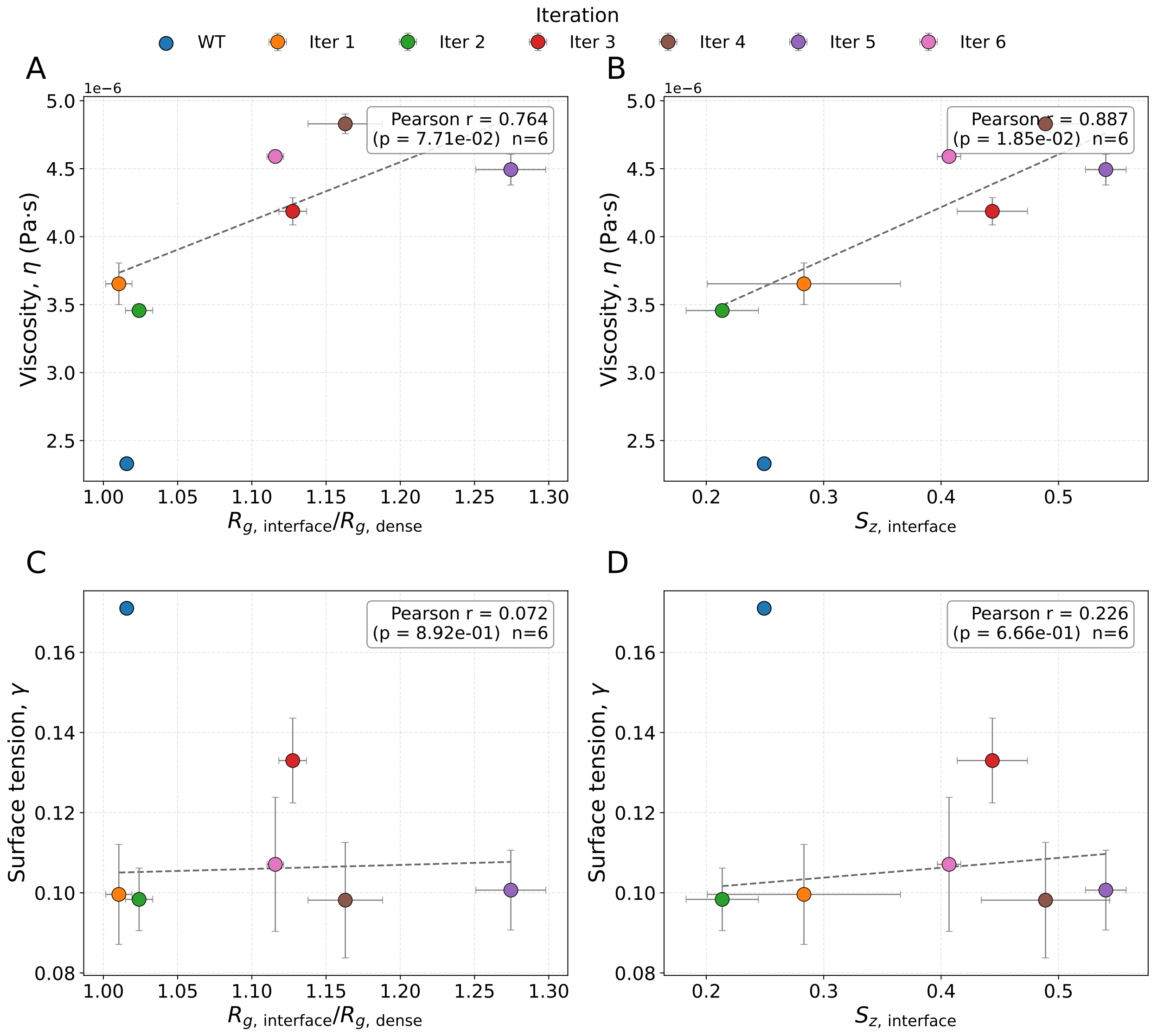


Figure S24: Correlations between the material properties of the MUT-16 condensate and the interfacial conformational properties of MUT-16 chains across MUT-8 mutant iterations with 50 peptides. **A.** Viscosity $\eta$ vs. $R_{g,\text{interface}}/R_{g,\text{dense}}$. **B.** $\eta$ vs. $S_{z,\text{interface}}$. **C.** Surface tension $\gamma$ vs. $R_{g,\text{interface}}/R_{g,\text{dense}}$. **D.** $\gamma$ vs. $S_{z,\text{interface}}$.
